# A comprehensive resource for studying microRNA evolution and microRNA-mediated development and whole-body regeneration in the acoel worm *Hofstenia miamia*

**DOI:** 10.1101/2024.12.01.626237

**Authors:** Ye Duan, Tomer Segev, Danielle E. Khost, Timothy B. Sackton, Isana Veksler-Lublinsky, Victor Ambros, Mansi Srivastava

**Affiliations:** Program in Molecular Medicine, UMass Chan Medical School, Worcester, MA, USA; Department of Organismic and Evolutionary Biology, Museum of Comparative Zoology, Harvard University, Cambridge, MA, USA; The Stein Faculty of Computer and Information Science, Ben-Gurion University of the Negev, Beer-Sheva, Israel; FAS Informatics Group, Harvard University, Cambridge, MA, USA

**Keywords:** microRNA, *Hofstenia miamia*, acoel, xenacoelomorph, pri-microRNA, spliced intron, miRNA targets

## Abstract

The acoel worm *Hofstenia miamia* has recently emerged as a model organism for studying whole-body regeneration and embryonic development, providing an opportunity to probe post-transcriptional mechanisms in both processes in the same organism. Here, we establish a resource for studying *H. miamia* microRNA-mediated gene regulation, a major aspect of post-transcriptional control in animals. We generated a PacBio long-read sequencing-based genome assembly. We developed a stringent microRNA annotation framework for *H. miamia* using small RNA-sequencing samples spanning key developmental stages. Our analysis uncovered 545 microRNA loci, including 154 highest-confidence loci based on structural features, expression levels, and prediction quality metrics. Comparison of microRNA seed sequences with those in other bilaterian species revealed that *H. miamia* encodes many known conserved bilaterian microRNA families and that several microRNA families previously reported only in protostomes or deuterostomes likely have ancient bilaterian origins. We profiled and characterized the expression dynamics and strand preference of microRNAs in *H. miamia* embryonic and post-embryonic development. An intron that is spliced in the primary transcript of co-transcribed *let-7* and *mir-125* microRNAs. To generate hypotheses for microRNA function, we annotated the 3’ UTRs of *H. miamia* protein-coding genes and performed microRNA target site predictions. Focusing on genes that are known to function in the wound response, posterior patterning, and neural differentiation in *H. miamia*, we found that these processes may be under substantial microRNA regulation. Notably, we found that microRNAs in MIR-7 and MIR-9 families, which have target sites in the posterior genes *fz-1*, *wnt-3*, and *sp5* are indeed expressed in the anterior of the animal, consistent with an anterior-biased repressive effect on their corresponding target genes. Our annotation provides a resource for future studies of post-transcriptional regulation of gene expression during development and regeneration.

## BACKGROUNDS

Embryonic development and regeneration both require dynamic and rapid regulation of gene expression. While both processes have been investigated extensively at the level of transcriptional regulation, the role of post-transcriptional regulation is relatively less understood, particularly during regeneration. MicroRNAs (miRNA) are endogenous, non-coding RNAs present in multi-cellular organisms that play crucial roles in post-transcriptional gene regulation, particularly in biological processes that require precise control of complex and dynamic gene expression patterns (1, 2). miRNAs are encoded by specific gene loci and transcribed into primary transcripts (pri-miRNA), which are processed into precursor miRNAs (pre-miRNA).

These pre-miRNAs consist of the miRNA strand and the passenger strand arranged in a hairpin structure (1, 3). The pre-miRNA is further processed into an RNA duplex where the two strands of the duplex are referred to as 5p and 3p strands (1). Within this complex, one strand is loaded into the Argonaute protein while the other strand is degraded (1, 3). The miRNA-Argonaute complex subsequently binds to the 3’ untranslated region (UTR) of target messenger RNAs (mRNAs) by partial complementary base-pairing and represses the protein production (4).

Given the likely extensive influence of miRNAs in regulating gene expression, a comprehensive understanding of animal biology warrants investigations of miRNAs for metazoan biological processes such as embryogenesis and regeneration.

A full understanding of the roles of miRNAs in animal biological processes also requires a study of miRNA evolution. miRNAs are grouped into seed families based on identity of the seed sequence, corresponding to nucleotides 2-8 from the 5’ end (g2-g8), which reflects the potential for recognition of common targets (5). Further, miRNAs are also categorized into gene families based on the inferred phylogeny of their pre-miRNA sequences and genomic synteny (6, 7).

Studies of metazoan genomes have revealed that certain miRNAs, for example, those in the MIR-100 family, originated from an eumetazoan ancestor (which gave rise to cnidarians and bilaterally symmetric animals, bilaterians), whereas other miRNAs, for example the LET-7 family, originated at the base of bilaterians 550 million years ago (8). Although miRNAs have been annotated in many bilaterian species, key animal lineages that could inform the origins of bilaterian miRNAs have very limited data available. Only 121 miRNAs have been reported thus far from acoels, nemertodermatids, and xenoturbellids, which together compose the phylum *Xenacoelomorpha*, potentially the earliest diverging bilaterian lineage (Figure 1A) (9–12). We therefore sought to expand the catalog of xenacoelomorph miRNAs.

**Figure 1.**
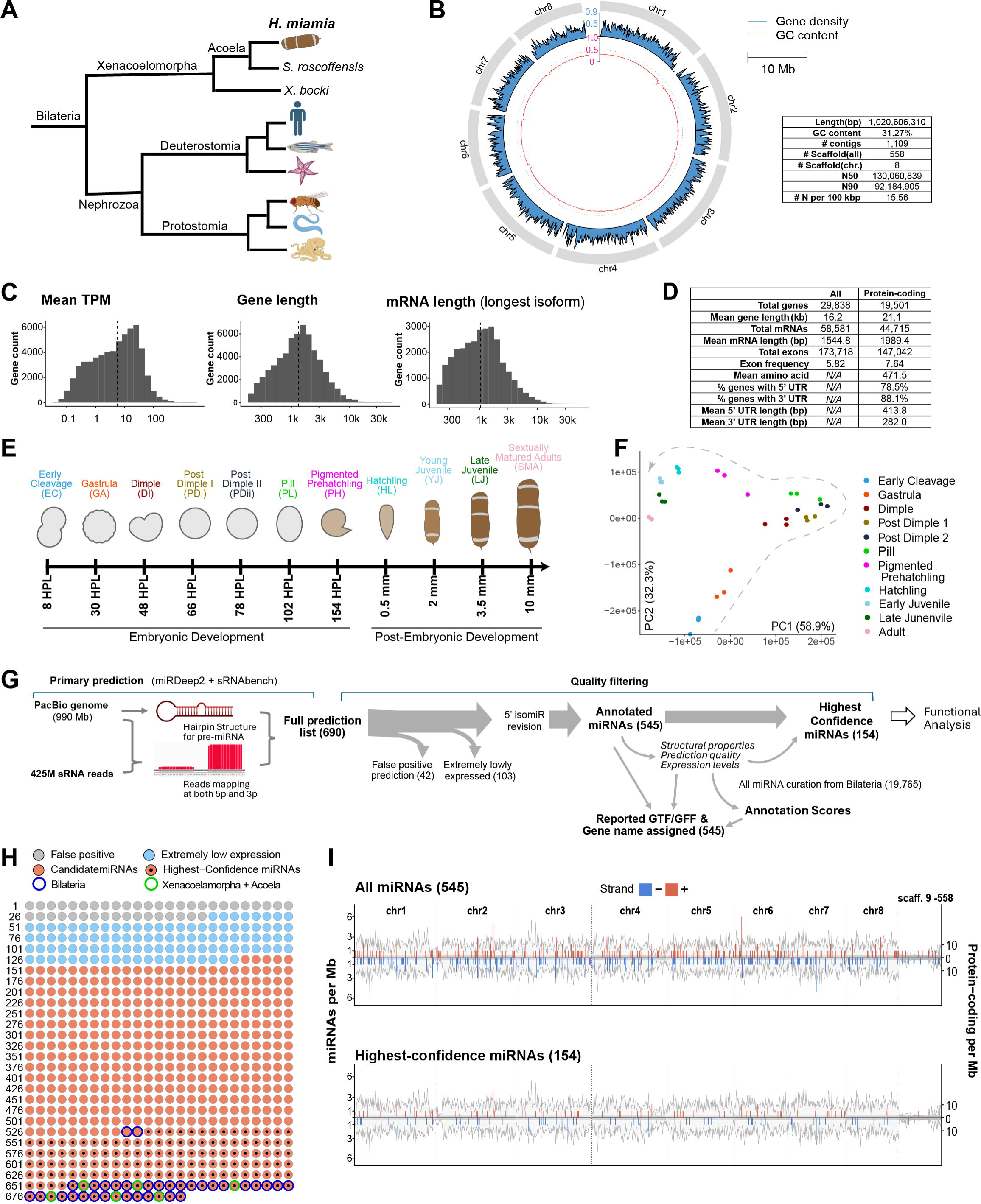
PacBio genome assembly and miRNA annotation in *H. miamia*. **A.** Phylogenetic tree showing the evolutionary position of *H. miamia* among other bilaterian species. **B.** Circos plot of the PacBio long-read genome assembly showing chromosome length, gene density and GC content. **C.** Summary statistics of the genome annotation, including the distributions of mean TPM (left), gene length (middle), and length of the longest mRNA for each gene (right). Vertical dashed line indicates the mean value. **D.** Statistical table for the genome annotation. **E.** Schematic of the embryonic and post-embryonic developmental stages sampled for miRNA annotation. HPL, hours post-egg laying. **F.** Principal component analysis (PCA) of normalized miRNA count across developmental stages. Each developmental stage includes three biological replicates. The dashed curve indicates the developmental trajectory connecting sequential stages. **G.** Overview of the miRNA annotation pipeline. **H.** Filtering results and analysis of the seed family of all the 690 predictions. X/A denotes Xenacoelomorpha/Acoela. **I.** Genomic distribution of all 545 annotated miRNAs (top) and the 154 highest-confidence miRNAs (bottom). miRNA density was calculated using 1 Mbp bins, with gene density shown in the grey background.

We focused on the acoel *Hofstenia miamia,* also known as the three-banded panther worm, a marine invertebrate that has recently emerged as a favorable research organism for studying regeneration and embryonic development (13–16). This is largely due to its robust capacity for whole-body regeneration in both juvenile and adult stages, and for producing embryos suitable for morphological and genetic studies. Notably, unlike planarians, another well-established system for whole-body regeneration, the accessibility of *H. miamia* embryos can be utilized for stable transgenesis and CRISPR/Cas9 genome editing (17). Also, *H. miamia* can undergo both sexual and self-reproduction, which enables genetic crossing. These features make *H. miamia* a promising system for genetic studies of miRNAs in regeneration and embryogenesis.

Current research in transcriptomics and chromatin accessibility has revealed an extensive catalog of dynamic transcriptional regulation during *H. miamia* whole-body regeneration (18). However, there are still phenomena that cannot be explained by transcriptional regulation alone. For example, the *H. miamia* anterior-posterior (AP) axis is regulated by the expression of *Wnt* ligand *wnt-3*, whose mRNA level is upregulated within 3 hours post-amputation specifically at posterior-facing wounds, indicating the re-establishment of pattern along the AP axis (19).

However, the expression of the wound-induced transcription factor *egr*, which is needed for wound-induced *wnt-3* expression, as well as increased chromatin accessibility at the *wnt-3* locus, occurs symmetrically at both anterior- and posterior-facing wound sites (18, 19).

Therefore, post-transcriptional regulation, potentially by miRNAs, must be considered as a plausible explanation for asymmetric upregulation of *wnt-3* mRNA.

In this study, we present a comprehensive annotation of *H. miamia* miRNAs along with developmental profiling of their expression. We provide a new chromosome-level genome assembly for *H. miamia* using PacBio long-read sequencing, with which we annotated 545 miRNA loci, including a subset of 154 highest-confidence miRNAs. Compared to previous *H. miamia* miRNA annotations, our work incorporated in-depth sequencing data and utilized comprehensive sampling of embryonic and post-embryonic development stages (12, 20). We also developed a scoring system for the nominated miRNA loci based on predicted biophysical properties and expression parameters of candidate pre-miRNAs using computational tools and the set of curated bilaterian miRNAs as a reference. By analyzing the seed families of the annotated *H. miamia* miRNAs, we demonstrate that our annotation provides candidates for revising the current understanding of the evolutionary origins of certain miRNA families.

Additionally, we analyzed the expression patterns of miRNAs across developmental stages and observed several notable phenomena. For example, the conserved *let-7* and *mir-125* miRNAs were unconventionally enriched in the early embryonic stages due to maternal deposition, and some miRNAs, including *mir-124,* exhibited developmentally regulated 5p/3p strand preference alterations. We also identified a spliced intron within the primary transcript of the clustered *let-7* and *mir-125* miRNAs. To investigate the potential target repertoire of the *H. miamia* miRNAs, we performed StringTie annotation of *H. miamia* expressed mRNAs with integration of our previous transcriptome and performed computational miRNA target predictions using the 3’ UTR sequences (21). Our results reveal that miRNAs in certain seed families show enrichment for target sites within the gene groups that have been genetically characterized for specific functions in regeneration and development. Notably, some miRNAs exhibited anterior enriched expression patterns, which anti-correlated with the expression pattern of key posterior- expressed protein-coding genes. This work can serve as a resource for future studies of the functions of miRNAs and their targets, and the regulation of microRNA expression, in *H. miamia* development and regeneration.

## RESULTS

### PacBio-based genome assembly shows high continuity

Previously, we published a *H. miamia* genome assembled using short-read sequencing and scaffolded by Hi-C and Nanopore sequencing (18). To further enhance genome completeness for miRNA prediction, we performed a *de novo* genome assembly using PacBio long-read sequencing (hereafter referred to as the PacBio genome). The PacBio assembly yielded a 1.02 Gb genome comprising 8 chromosomes and 550 small scaffolds, with a N50 of 130M (Figure 1B). Notably, this new assembly reduces gapped regions by 73%, to 15.6 Ns per 100 kbp compared to 56.59 Ns per 100 kbp in our previous short-read assembly, indicating a substantial improvement in genome completeness and contiguity.

To provide the genomic and transcriptomic resources required for downstream analyses of *H. miamia* miRNAs, we generated a *de novo* protein-coding gene annotation of the PacBio genome using pair-end RNAseq datasets with multiple developmental samples (13, 22). We further improved the annotation by incorporating a recently published transcriptome assembly followed by manual curation (22). The final annotation contained 29,838 gene loci, including 19,501 protein-coding genes, and achieved 93% complete BUSCOs against Eukaryota_OBD12 dataset (Figure 1C-D, S1C).

### Prediction of *H. miamia* miRNA loci

To annotate the *H. miamia* miRNA complement, we performed small RNA sequencing (sRNA-seq) using total RNA samples across the *H. miamia* life cycle. Previous studies in various animals have shown that miRNA expression can be highly dynamic during different developmental stages (23–25). Therefore, to ensure comprehensive identification of miRNAs, particularly those that are developmentally regulated, we collected total RNA samples from seven embryonic and four post-embryonic developmental stages (Figure 1E-F, S2A-B). These time points were selected to cover the developmental stages undergoing the major transcriptome changes as previously described, where miRNA-mediated regulation could play a significant role (14). After quality filtering and removal of ribosomal RNA reads, we obtained in total 425 million sRNA reads, with an overall mapping rate of 83% to the PacBio genome. We then used these sRNA reads to predict miRNA loci in the *H. miamia* genome using miRDeep2 and sRNAbench (26, 27). Broadly, both algorithms identify miRNA precursors by scanning for potential stem-loop structures in the genome and defining a pre-miRNA locus if reads can be mapped to both the 5p and 3p arms of the pre-miRNA (Figure 1G).

Using default settings, we identified 690 potential miRNA loci, with 554 loci predicted by miRDeep2 and 278 predicted by sRNAbench, of which 142 were predicted by both algorithms (Figure 1H). Of these, we found 42 loci lacking plausible pre-miRNA hairpin structures and flagged these predictions as false positives (Figure 1H). These include loci that (a) failed to fold into any secondary structure (n=17); (b) exhibited insufficient base-pairing in the stem region to support a stable hairpin structure (n=16); (c) displayed secondary structures with noncanonical pre-miRNA hairpins (n=8; Figure S2C); or (d) formed a predicted chimeric hairpin assembled from strands of two nearby pre-miRNAs (n= 1; Figure S2D). Because some miRNAs share mature or pre-miRNA sequences that can confound locus-level counts in miRDeep2 and sRNAbench, we quantified the miRNA expression of the retained loci using fractional counting (Figure S2E). 103 loci were also flagged as false positives due to extremely low expression.

These included 83 loci whose mature or star strand had less than one cumulative fractional reads across all 33 libraries; 8 loci for which no single library produced more than one fractional read in any of the 33 libraries; and 12 loci with in-cluster ratio less than 50%, suggesting that most reads were not derived from the mature or star strand (Figure 1H). Note that, because we later observed that some miRNAs express both 3p and 5p strands and that the dominant strand can switch during development (Figure 4E-H), we evaluated each locus using the in-cluster ratio considering both the mature and star strands, rather than the ratio of the mature strand alone (Figure S3) (28). Some miRNA loci exhibited heterogeneity at the 5’ end of the expressed mature miRNA; for the 22 miRNA loci that exhibited >50% 5’ end heterogeneity, we annotated their 5’ coordinates and sequences, accordingly, and re-evaluated the predicted structural features of the corresponding pre-miRNAs (see Material and Methods; Figure S4).

### Definition of the highest-confidence miRNA subset

After removing the false-positive predictions, 545 miRNA loci were retained, including 426 loci predicted by miRDeep2 and 259 predicted by sRNAbench, with140 that had been predicted by both algorithms. We refer to these as the nominated *H. miamia* miRNAs and incorporated them into the genome annotation (Figure 1I). However, some loci still exhibited suboptimal predicted pre-miRNA structures or low read abundance. Although these loci could not be confidently classified as false positives, they arguably represented lower-confidence predictions for downstream analyses. Correspondingly, we generated a subset of highest-confidence miRNA loci prioritized for our subsequent study. A miRNA was included in the highest confidence subset only if it met all the following criteria: (a) The length of the 5p and 3p strands ranged between 20-26 nucleotides as recommended by MirGeneDB; (b) the minimum free energy (MFE), mature base-pairing ratio, and loop size of the pre-miRNA fell within the 99^th^ percentile range of all pre-miRNAs curated in MiRGeneDB 3.0 across available bilaterian species; (c) the maximum bulge asymmetry was less than 3 nt to avoid major helix distortion (29); (d) the 3’ overhang was between 0-3 nt, consistent with Dicer processing (30–32); (e) the combined mature and star strand fractional read count exceeded the cumulative-abundance threshold corresponding to the top 99.9% of all mapped reads (1995 reads); (f) the maximum mature per-base coverage is above 50th percentile among *H. miamia* miRNAs in this study; and (g) 5’ heterogeneity ≤20% and the mature-plus-star in-cluster ratio ≥90%, based on empirically defined thresholds. This highest-confidence subset includes a total of 154 miRNA loci, 121 of which had been identified by miRDeep2, 117 by sRNAbench, with 84 loci identified by both (Figure 1H-I, Table S2).

### Scoring system for candidate miRNAs

To estimate the confidence of assigning miRNA status to each candidate miRNA locus without arbitrary false negative and false positive cutoffs, we developed a scoring system based on structural features, read abundance, and prediction-quality metrics (see Methods, Figure 2A). Structural penalties were assigned according to the mature and star strand lengths, and predicted structural parameters of the pre-miRNA, including minimum free energy, mature-strand base-pairing ratio, apical loop sizes, maximum bulge asymmetry, and 3’ overhang (Figure 2B-G and S5A). Scoring penalty thresholds for these features were determined using the pre-miRNAs curated in MirGeneDB 3.0 from all available bilaterian species (6, 12, 29) (Table S1).

**Figure 2.**
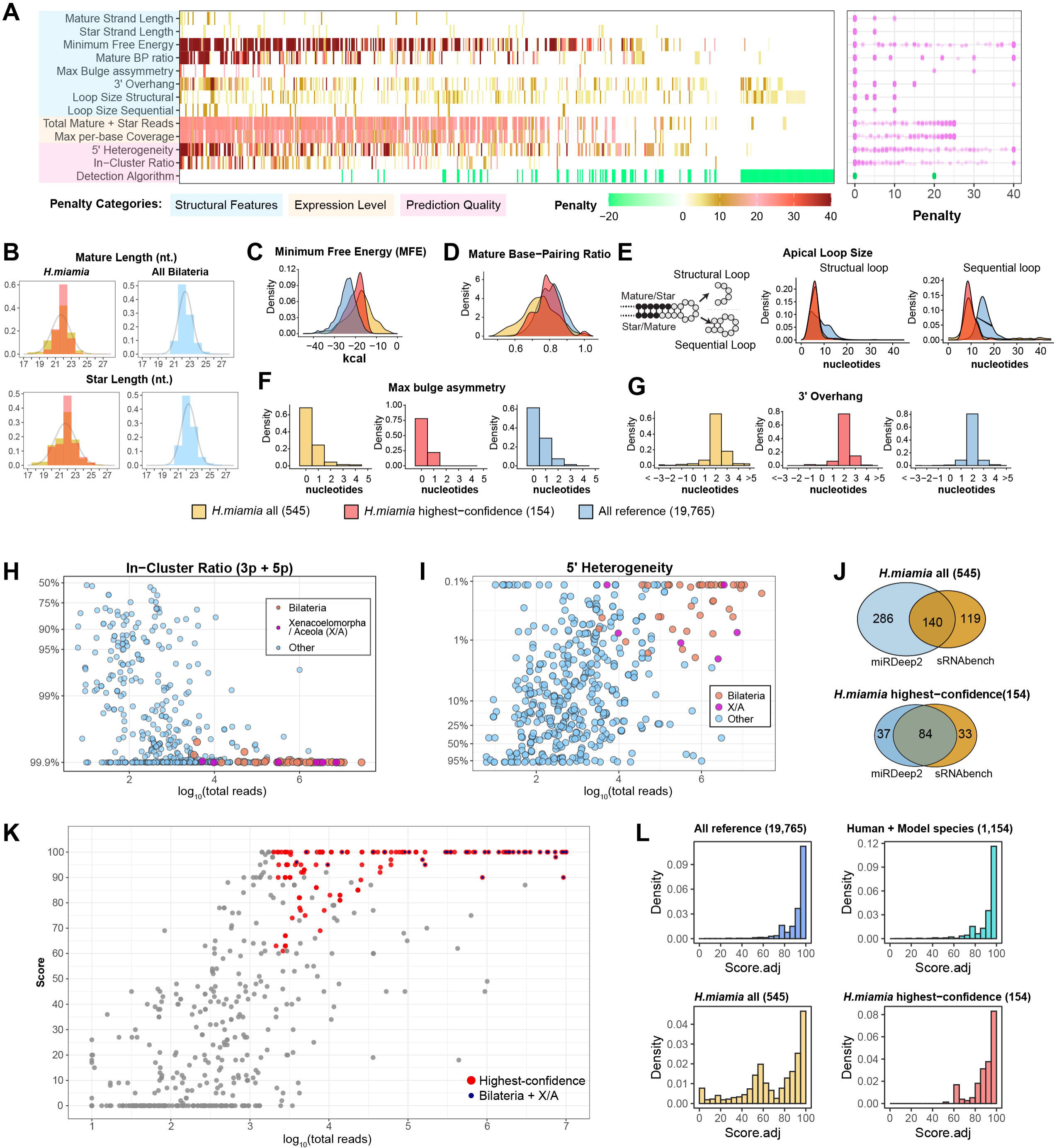
Structural features, prediction quality, and scoring system of *H. miamia* miRNAs. **A.** Summary of the scoring system for the nominated *H. miamia* miRNAs. The heatmap (left) shows the penalty assigned to each miRNA (columns) per feature (rows). Detection algorithms are represented as negative penalty values. The dot plot (right) shows the distribution of penalty (magenta) or bonus (green) per evaluated feature. **B-G.** Distribution of structural features for all and the highest-confidence *H. miamia* miRNAs, and all curated bilaterian miRNAs including mature/star strand length (B), minimum free energy (C), mature base-pairing ratio (D), structural and sequential apical loop size (E), maximum bulge asymmetry (F), and 3’ overhang (G). Negative values of the 3’ overhang indicate 5’ overhang. **H.** Scatter plot of in-cluster ratio in percentage (y-axis) against the total read counts (x-axis). **I.** Scatter plot of the - log_10_ of 5’ heterogeneity (y-axis) against the total read counts (x-axis). The 5’ heterogeneity was calculated after sequence revision (Figure S4). **J.** Venn diagrams showing the detecting algorithm of all and highest-confidence *H. miamia* miRNAs. **K.** Scatter plot of the annotation scores (y-axis) against total reads (x-axis). Red points, highest-confidence miRNAs. Black inlays, miRNA whose seed families are assigned to an evolutionary origin of Bilateria or Xenacoelomorpha/Acoela. **L.** Distribution of adjusted annotation scores for all and the highest-confidence *H. miamia* miRNAs, all curated Bilaterian miRNAs, and curated miRNAs from human and four conventional model organisms (*C. elegans, D. melanogaster, D. rerio, M. musculus*) in MirGeneDB 3.0. The adjusted annotation score was calculated by excluding the penalties for read counts, in-cluster ratio, 5’ heterogeneity, and detection algorithms.

Expression-based penalties were assigned according to the cumulative abundance of mature and star strands and the maximum mature-strand per-base coverage (Figure S5B-C; Table S1). Prediction-quality penalties were assigned based on combined 3p/5p in-cluster ratio and 5’ heterogeneity, and whether the locus was predicted by both algorithms (Figure 2H-J and S5D-F). The final annotation score was calculated by combining these penalties as described in Methods. The feature values, penalties, and annotation scores are provided in the output GTF file (Supplemental Dataset S2, Table S3).

Compared to the complete set of 545 nominated miRNA loci, or to the excluded (non-highest-confidence) subset, the set of 154 highest-confidence *H. miamia* miRNAs exhibited significant enrichment for both high scores and high read count without significant change of the nucleotide preferences (Figure 2K; Figure S6A, S6C). The number of highest-confidence *H. miamia* miRNAs (154) is consistent with the pattern of correlation between genome size and number of annotated miRNA loci for several representative bilaterian species with high-quality miRNA annotations including human and conventional model organisms (Figure S6B).

To independently evaluate the scoring system and the highest-confidence miRNA subset, we calculated an adjusted score for miRNAs from all curated bilaterian species, including human and conventional model organisms, as described in Methods. We found that the complete set of 545 putative *H. miamia* miRNA loci contained a marked enrichment of low-scoring loci relative to the reference miRNAs, whereas this low-score tail was markedly reduced in the highest-confidence miRNA subset (Figure 2L). This independent comparison further supports the conclusion that the stringent filtering substantially reduced the rate of potentially false-positive annotations. We used this highest-confidence miRNA subset for all subsequent analyses in this study.

### Nomenclature of *H. miamia* miRNAs

We named each of the 545 nominated *H. miamia* miRNA loci based on the identity of their seed sequence, following established miRNA naming guidelines (33). To name the predicted miRNAs, we first extracted the g2-g8 7mer seed sequences from both the 5p and 3p strands of each pre-miRNA locus. For each seed, we searched for identical 7mer seed sequences among all mature miRNAs curated in MirGeneDB 3.0 from all bilaterian species and assigned a *H. miamia* seed name based on the family classifications in MirGeneDB 3.0. When the seeds of both the 5p and 3p strands of a *H. miamia* pre-miRNA matched MirGeneDB 3.0 families, the pre-miRNA was named according to the more abundantly expressed strand. Notably, in MirGeneDB, “ACCCUGU” (commonly known as the MIR-10 family), “CCCUGAG” (commonly known as the MIR-125/LIN-4 family), and “ACCCGUA” (commonly known as the MIR-100 family) are all classified and named as MIR-10 family due to their apparent common evolutionary origin (6).

However, genetic and biochemical studies show that these three families of miRNAs can have varied expression patterns and distinct biological functions in various species (2, 34–36), consistent with their distinct seed sequences. In this study, we assigned and named the “ACCCUGU” seed as the MIR-10 seed family, and the “CCCUGAG” as the MIR-125 seed family (Figure S7H). We did not detect the “ACCCGUA” seed in *H. miamia*, but we refer to this family as MIR-100 in our subsequent conservation analyses (Figure S7H). The *H. miamia* g2-g8 seeds without any 7mer matches in MirGeneDB 3.0 were assigned names in the format “HMI-NOVEL- [number]”, where the numbering was determined by ranking the seed families in descending order based on total read counts. All *H. miamia* pre-miRNAs are prefixed with “Hmi-” to indicate the species. Loci expressing miRNAs with identical seed sequences but different non-seed sequences were distinguished by adding an “-[alphabet]” suffix to the family name, and those expressing miRNAs with identical mature sequences were further distinguished by a “-[number]” suffix. For example, Hmi-miR-124-b-1 and Hmi-miR-124-b-2 share both identical seed and non-seed sequences, whereas Hmi-miR-124-a shares identical seed sequence with Hmi-miR-124-b- 1/2 but differs in the non-seed region (Figure 3A). Note that in MirGeneDB, a curated species-specific family is named using the format of “SPECIES-NOVEL-”, which distinguishes it from the “HMI-NOVEL-” format. For example, the seed “UUCUCAU” had a 7mer match to the previously *Canis familiaris*-specific family CFA-NOVEL-13, so the *H. miamia* miRNA with the seed “UUCUCAU” were named as hmi-cfa-novel-13. This “HMI-CFA-NOVEL-1” seed family is distinct from the seed family “HMI-NOVEL-13”, which is specific to *H. miamia* with a different sequence of “UUCCAUU”. Also, we use all capitalized letters to indicate the seed family (e.g., MIR-7 family), italicized lower-case letters to indicate miRNA genes (e.g., *hmi-mir-1*), and a hybrid format to indicate the miRNA molecule (e.g., Hmi-miR-1)(33).

**Figure 3.**
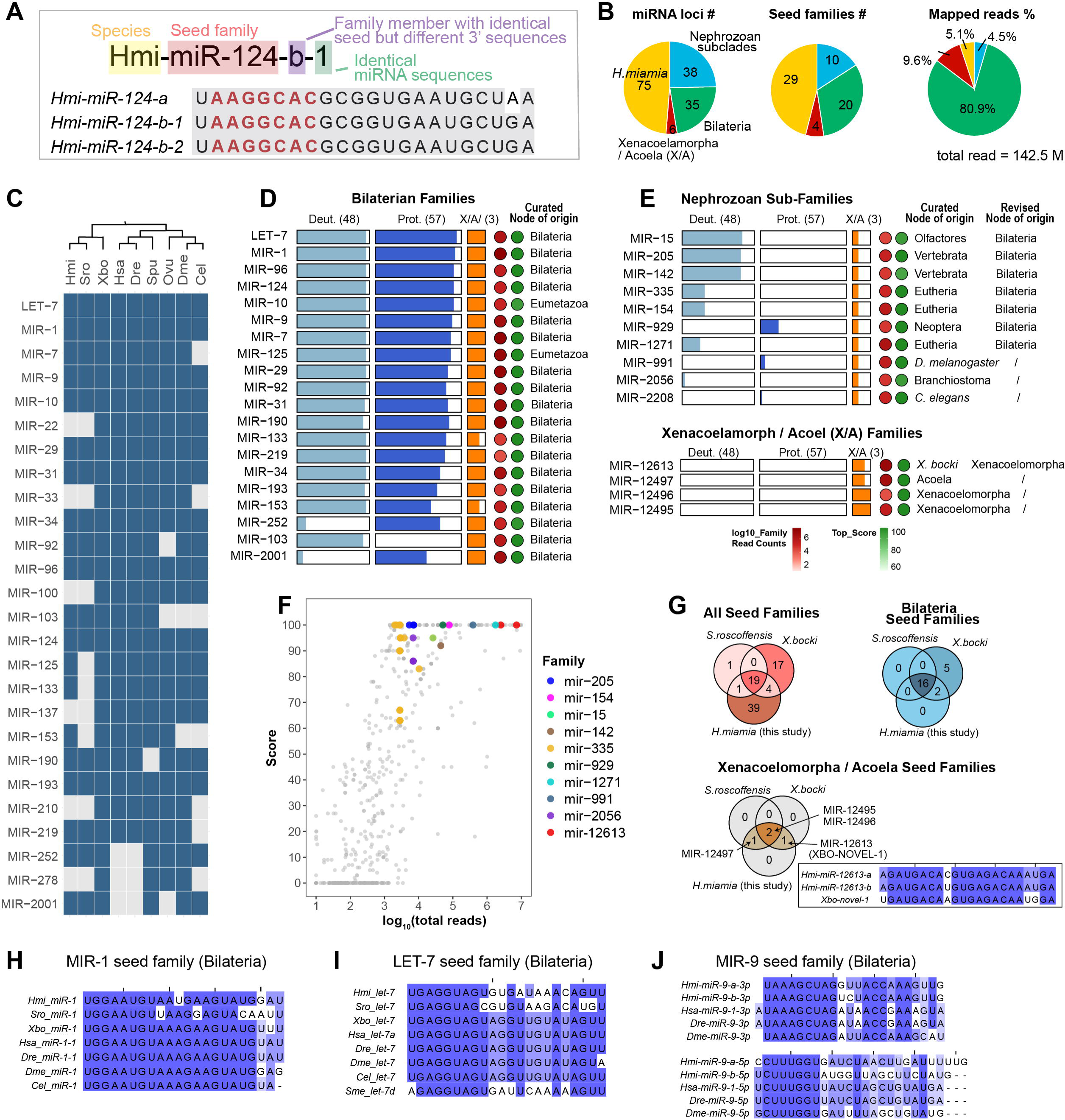
Evolutionary analysis for *H. miamia* seed families. **A.** Illustration of *H. miamia* miRNA naming guidelines using the example of hmi-miR-124-b-1. **B.** Pie charts showing the numbers of miRNA loci (left), numbers of seed families (middle), and distribution of total mapped miRNA read counts categorized by the evolutionary node of origin. **C.** Presence or absence of all bilaterian seed families in representative bilaterian species including *H. miamia* (Hmi)*, Symsagittifera roscoffensis* (Sro)*, Xenoturbella bocki* (Xbo), Homo Sapiens (Hsa), *Derio rerio (Dre), Strongylocentrotus purpuratus* (Spu), *Octopus vulgaris* (Ovu), *Drosophila melanogaster* (Dme), and *Caenorhabditis elegans* (Cle). **D-E.** Occurrence of identified bilaterian seed families (**D**) and nephrozoan sub-clades or xenacoelomorph/ acoel family (**E**) across curated species with miRNA annotation. The bars represent the proportion of species within corresponding evolutionary groups for each seed family. The color of the red points indicates the summed total read counts of all miRNAs within the seed family. The color of the green points indicates the highest score among miRNAs within the seed family. The node of family origin was annotated based on MirGeneDB 3.0 (12). **F.** Scatter plot of the annotation scores against expression, highlighting miRNAs whose curated nodes of origin are currently assigned to nephrozoan sub-clades or *X. bocki* but identified in *H. miamia* (Figure 3G). **G** Venn diagrams of the numbers of all seed families (top left), seed families annotated as bilaterian/eumetazoan (top right), and seed families annotated as xenacoelomorph/acoel (bottom) in *Hmi, Sro, and Xbo.* Inset shows the sequence alignment of XBO- NOVEL-1 family miRNAs in *X. bocki* and *H. miamia*. **H-J.** Sequence alignments of MIR-1 (**H**), LET-7 (**I**), and MIR-9 (**J**) families. Sequence alignments and conservation were visualized using JalView (104). All analyses in this figure were performed using the 154 highest-confidence miRNAs as input.

### Evolutionary origins of miRNA seed families identified in *H. miamia*

We classified the seed families of the 154 highest-confidence *H. miamia* miRNAs using the curated miRNA seeds from all metazoan species in MirGeneDB 3.0 as references. We found 39 unique 7-mer seed sequences from 79 miRNA loci that matched curated miRNA seed sequences. According to the MirGeneDB classification, these 39 distinct seed sequences correspond to 34 seed families, as in certain cases, multiple seed sequences have been assigned to the same family. For example, “AUUGCAC” and “AUCGCAC” are both assigned to the MIR-92 family, in this case, based on the potential for G:U wobble base pairing at position g4 of “AUUGCAC” to the same target sequence as by “AUCGCAC”. In addition, we identified 29 unique seed sequences from 75 miRNA loci that did not match any curated miRNA seeds and were therefore assigned to 29 *H. miamia-*specific seed families. Among the 63 seed families in *H. miamia* (34 conserved seed families plus 29 thus far *H. miamia* specific seed families), 29 families are encoded by multiple miRNA loci, with the largest family, MIR-335, comprising 21 miRNA loci (Figure S7A). We also identified 22 groups of miRNA loci with identical mature sequences (84 loci in total) across 19 families, and 18 groups with identical pre-miRNA sequences (21 loci in total) across 15 families (Figure S7B). Notably, MirGeneDB 3.0 had previously curated 20 *H. miamia* seed families from 27 miRNAs, and they were all recovered in our annotation (Figure S7C).

We next analyzed the seed families of the highest-confidence *H. miamia* miRNAs using the curated nodes of family origin in MirGeneDB 3.0 (Figure 3B, Table S4). MirGeneDB 3.0 currently curates 25 bilaterian seed families as bilaterian, based on their presence in both *Xenacoelomorpha* and *Nephrozoa* (Figure S7F). The *Xenacoelomorpha* phylum includes *H*.

*miamia* and *Symsagittifera roscoffensis (Sro)* from the *Acoela* clade, as well as *Xenoturbella bocki (Xbo)* which is sister to *Acoela* (Figure 1A). We identified 20 bilaterian seed families from 35 miRNA loci in *H.* miamia, and these accounted for 80.9% of all mapped miRNA reads (Figure 3B-D, S7E-F). There were five bilaterian families (MIR-22, MIR-33, MIR-137, MIR-210, MIR-278) not found in *H. miamia* were undetected in *Sro* but detected in *Xbo* (Figure 3C-D, S2G).

Similarly, the eumetazoan family MIR-100 was not found in either *Sro* or *H. miamia* but was detected in *Xbo* (Figure 3C-D, S7F). These findings suggest that these six seed families may have been lost in acoels (represented by *Hmi* and *Sro*) but persist in other bilaterian species. Interestingly, three bilaterian seed families are found in both *H. miamia* and *Xbo* but not in *Sro* (MIR-125, MIR-133, MIR-153) (Figure 3C-D, S7F). This pattern may indicate either that these families were lost in the evolution of *Sro*, or that the absence of annotation in *Sro* was a false-negative due to lack of sufficient sequencing depth and sampling (see Discussion).

In addition to the seed families curated as bilaterian families, the set of 154 highest-confidence *H. miamia* miRNAs includes 10 seed families (corresponding to 38 miRNA loci) that are currently annotated as specific to nephrozoan subclades, i.e. bilaterian groups excluding xenacoelomorph (Figure 3B, 3E). In general, miRNA loci from these nephrozoan subclade families exhibited significantly lower read counts and scores than miRNA loci from bilaterian families (Figure S7D), suggesting that they are more difficult to detect in *Xbo, Sro* because of their lower expression and/or sub-optimal precursor structures. Nevertheless, every family contained at least one miRNA with either a high annotation score, high abundance, or both (Figure 3F), and seven families (MIR-15, MIR-205, MIR-142, MIR-335, MIR-154, MIR-929, MIR-1271) have been detected in multiple species (Figure 3E). Therefore, these families may have escaped previous detection because of limited sequencing depth and/or developmental stage sampling, rather than indicating true innovations in their clades. Our *H. miamia* annotation, supported by deep sRNA sequencing and thorough coverage of developmental stages, indicates that these seed families are strong candidates for ancient bilaterian miRNA seed families. Exceptions may be the MIR-991, MIR-2208, and MIR-2056 families, which are currently curated only from three species of the *D. melanogaster* subgroup, *C. elegans,* and two *Branchiostoma* (lancelets) species, respectively (Figure 3E). Their presence in *H. miamia*, together with their restricted phylogenetic distribution, suggests that these families more likely arose from convergent evolution than shared ancestry.

Further, we identified two *Xenacoelomorpha*-specific families, MIR-12495 and MIR-12496, as well as the *Acoela*-specific family MIR-12497 (Figure 3E-G). Notably, we found that the XBO-NOVEL-1 seed family is present in both *H. miamia* and *X. bocki,* with 11 out of 14 non-seed sequences identical (Figure 3G). In addition, the two *H. miamia* XBO-NOVEL-1 miRNAs both exhibited high expression levels and high annotation scores (Figure 3F). These findings indicate that this family, which is currently annotated as *Xbo*-specific, should instead be classified as *Xenacoelomorpha*-specific. Accordingly, we re-named this miRNA family as MIR-12613, following the MirGeneDB nomenclature principles for families identified in multiple species.

### *H. miamia* MIR-1/LET-7/MIR-9 miRNAs exhibit different patterns of conservation in the 3’ non-seed region

Besides the seed region, nucleotide sequences in the 3’ non-seed region (g9-g22) have also been reported to critically contribute to target repression and miRNA degradation (37–39).

Particularly, LET-7 and MIR-1 families exhibit high levels of conservation at the 3’ non-seed region (37). In *H. miamia*, we found that the hmi-miR-1 mature sequence is also conserved with only 2 nt different from the best pan-bilaterian consensus sequence (Figure 3H). Interestingly, the MIR-1 miRNAs exhibit variation of the 3’ distal sequence between *H. miamia, Sro,* and *Xbo* (g20-g22), despite the deep conservation in other non-seed regions (Figure 3H). This pattern, namely deep conservation with variation at the 3’ distal region, is also observed for the MIR-1 family in all bilaterians (37).

Like MIR-1, the LET-7 family also includes a miRNA whose sequence is nearly fully conserved at every non-seed position across most bilaterian species (8, 37). However, in *H. miamia* and *Sro*, we did not detect this deeply conserved LET-7 family member (Hsa-let-7a, referred to as let-7a in this paper). Instead, the identified Hmi-let-7 and Sro-let-7 miRNAs exhibit sequence variation in the g9-g18 region compared to the deeply conserved let-7a miRNA. In contrast, *Xbo* contains a LET-7 miRNA fully conserved with the *let-7a* (Figure 3I). It is noteworthy that despite the lack of conservation at positions g10-g18, the 3’ distal region (g19-g22) of Hmi-let-7 miRNA is identical to the deeply conserved *let-7a* (Figure 3I, see Discussion).

We also investigated the MIR-9 family, where both the 5p and 3p strands are evolutionarily conserved, and both strands can be expressed and contribute to biological functions in various species (40–43). In *H. miamia,* both MIR-9 family miRNAs possess the evolutionarily conserved MIR-9 seeds for both 5p (“CUUUGGU”) and 3p (“AAAGCUA”) (Figure 3J). Moreover, the 3p strands, which are predominantly expressed in *H. miamia* (Figure S7G), exhibited extensive conservation at positions g11-g19 (Figure 3J). The most abundant MIR-9 family member, hmi- miR-9-a, exhibited extensive non-seed sequence conservation for both 3p and 5p strands (Figure 3J). This conservation of both strands suggests that they may likely contribute to functional target repression in *H. miamia*.

### Developmental regulation of *H. miamia* miRNA expression

To investigate the expression dynamics of the identified miRNAs during *H. miamia* embryonic and post-embryonic development, we analyzed miRNA expression across eleven embryonic and post-embryonic stages (Table S5). The seven embryonic samples of early cleavage (EC), gastrula (GA), dimple (DI), post-dimple I (PDi), post-dimple II (PDii), pill (PL), and pigmented prehatchling (PH) were staged and described using the post-synchronization time (hours post-laying, HPL; Figure 1C). The four post-embryonic stages hatchlings (HL), young juveniles (YJ), late juveniles (LJ), and sexually mature adults (SMA) were staged and described using the size of the animal, as the hatched *H. miamia* animals can exhibit asynchronous growth (Figure 1C). We performed fuzzy c-means soft clustering of expression patterns for 154 highest-confidence miRNAs and obtained four distinct clusters (Figure 4A-B, Figure S8) (44, 45). Clusters 1 and 2 represent the miRNAs enriched during embryonic development, whereas Clusters 3 and 4 represent miRNAs enriched during post-embryonic development. Specifically, Cluster 1 miRNAs (n = 26) are predominantly enriched during early embryonic development before the dimple stage (48 HPL) when the *H. miamia* embryo is postulated to undergo maternal-zygote-transition and exhibit low expression levels in all subsequent stages (46). Cluster 2 miRNAs (n = 64) show an increase in expression after the dimple stage, followed by a decrease during later embryonic stages, and remain low in post-embryonic stages. Cluster 3 miRNAs (n = 21) are expressed at later embryonic stages, with enrichment at post-embryonic stages, but their expression levels decline around sexual maturation. Cluster 4 miRNAs (n = 43) begin to increase in expression during post-embryonic development, reaching peak expression at sexual maturation.

We next investigated whether miRNAs of different evolutionary origins exhibit distinct patterns of developmental dynamics. Across all developmental stages, miRNAs that originated in the bilaterian ancestor dominated in terms of total miRNA read proportions (Figure 4C-D), especially so for Cluster 1 (Figure 4C-D). Notably, we observed decreasing levels of *H. miamia-*specific miRNAs and an increasing proportion of xenacoelomorph/acoel miRNAs over developmental time, although both groups represented relatively small fractions of total miRNA reads across developmental stages (Figure 4C-D). This pattern suggests that the *H. miamia-*specific miRNA may be preferentially associated with early embryogenesis, whereas the xenacoelomorph/acoel miRNA may have greater roles in post-embryonic development (see Discussion).

### *let-7* and *mir-125-a* miRNAs are enriched in early embryonic development potentially due to maternal deposition

In *H. miamia,* we observed that Hmi-miR-125-a and Hmi-let-7, both functionally well-characterized in other model organisms, exhibited a consistent increase in expression during post-embryonic development, consistent with their expression patterns and functional contributions known in other species (2, 47–49). Interestingly, in addition to their enrichment in sexually mature adults (SMA), Hmi-let-7 and Hmi-miR-125-a also show substantial enrichment at early embryonic stages (Figure S9A). This contrasts with the observations in other species, where the let-7 and miR-125/lin-4 miRNAs typically exhibit very low expression during embryonic development (23, 47). To understand this unique pattern, we asked whether Hmi-let-7 and Hmi-miR-125-b miRNAs are actively transcribed during early embryonic development in *H. miamia* or alternatively are present as maternal deposits. We reasoned that if these miRNAs are newly expressed in early embryos, their genomic loci should exhibit high accessibility, indicating the transcription of the pri-miRNAs. To test this, we utilized previously described *H. miamia* chromatin accessibility datasets from embryonic ATAC-seq (46). Since the available *H. miamia* datasets lack chromatin accessibility information for later post-embryonic stages, we used the hatchling juvenile stage (HL) as a proxy for the post-embryonic development. We found that, compared to the post-embryonic stage, the putative promoter region of the *hmi-let-7* and *hmi- mir-125-a* cluster was significantly less accessible during early embryonic stages (Figure S9B).

To provide a contrast, we analyzed the chromatin accessibility of *hmi-mir-34-a*, which is exclusively enriched at early embryonic stages and shows no enrichment in post-embryonic stages. Unlike the *let-7/mir-125-a* cluster, the *hmi-mir-34-a* promoter exhibited significantly higher accessibility at early embryonic stages, with remarkably reduced accessibility at the post-embryonic stage (Figure S9C). This observation suggests that the unconventional enrichment of Hmi-let-7 and Hmi-miR-125-a miRNAs at early embryonic stages in *H. miamia* is potentially due to a maternal deposition rather than *de novo* transcription during embryogenesis.

### *H. miamia* miRNAs can exhibit altered 5p/3p strand preference during development

During miRNA biogenesis, the 5p and 3p arms of the pre-miRNA are processed into the 5p-3p duplex, and one strand (referred to as mature/guide strand) is loaded in the miRISC and functions in target repression, and the counterpart strand (referred to as star/passenger strand) is in most cases, ejected and degraded (1, 50). Most miRNAs have a strong selection preference between 5p or 3p strands, whereas in some cases, e.g. Hsa-miR-9, both the 5p and 3p, are expressed and each contributes to biological functions (40). Strand preference can also be regulated by cell type, pathology, or genetic alterations of miRISC functions (50–52).

To investigate whether *H. miamia* miRNAs exhibit developmentally dynamic 5p/3p strand preference, we analyzed the normalized expression of both the 5p and 3p arms and calculated the 5p/3p dominance in the context of development (Figure 4E-F). We defined strand preferences as where one strand is expressed at more than twice the level of the other (Figure S10A). Notably, for low-abundance miRNAs or developmental stages when both 5p and 3p strands are expressed at low levels, the accuracy of the 5p/3p ratio may be compromised as small changes or noise in these low-expression values can disproportionately affect the computed ratio. To address this limitation, we excluded developmental stages where both 5p and 3p strands are expressed below 1 RPM.

Among the highest-confidence *H. miamia* miRNAs, we found that 54 miRNAs exhibited at least one developmental stage without significant strand preference as described above. Among these, 20 miRNAs displayed a 5p/3p switch in strand preference (Figure 4F). For instance, *hmi- mir-92-a* showed high expression of both the 5p and 3p strands with no significant preference at 10 out of the 11 tested developmental stages (Figure 4G). Notably, its strand preference switched from 3p dominance during early cleavage stage to 5p dominance at the later embryonic and post-embryonic stages (Figure 4G). This phenomenon of dual strand expression and altered strand preference was also reported for MIR-92 miRNAs in *Drosophila melanogaster* (51). Another example was *hmi-mir-124-b-1* and *hmi-mir-124-b-2*, for which the 5p strand dominated during embryonic development, while the 3p strand dominated during post-embryonic development, suggesting that strand preferences could be developmentally regulated (Figure 4H). Meanwhile, although some miRNAs did exhibit a significant strand preference, the expression level of the less-dominant strand can substantially increase at specific developmental stages without strict correlation with the dominant strand, as for example, *hmi-mir-9-a/b* (Figure S10B).

### Clustered *let-7* and *mir-125-a* miRNAs are spliced in the primary transcript

We next examined the genomic organization of the identified *H. miamia* miRNAs. Among all 545 nominated miRNA loci, 109 miRNAs were organized into 50 genomic clusters (defined as groups of miRNAs located within 10 kb intervals) (Figure S11A). Among the highest-confidence miRNAs, 26 miRNAs were organized into 12 clusters (Figure 5A). To investigate whether these clustered highest-confidence miRNAs are likely co-transcribed, we examined chromatin accessibility using published bulk ATAC-seq datasets spanning embryonic development and late-juvenile stage (18, 46) (Figure S10B-C). Based on the chromatin accessibility of putative promoter regions, we inferred that 17 miRNAs from eight clusters are likely co-transcribed from shared putative promoters (Figure S11B), whereas the remaining clustered miRNAs either have independent putative promoters or are located within introns (Figure S11C).

**Figure 4.**
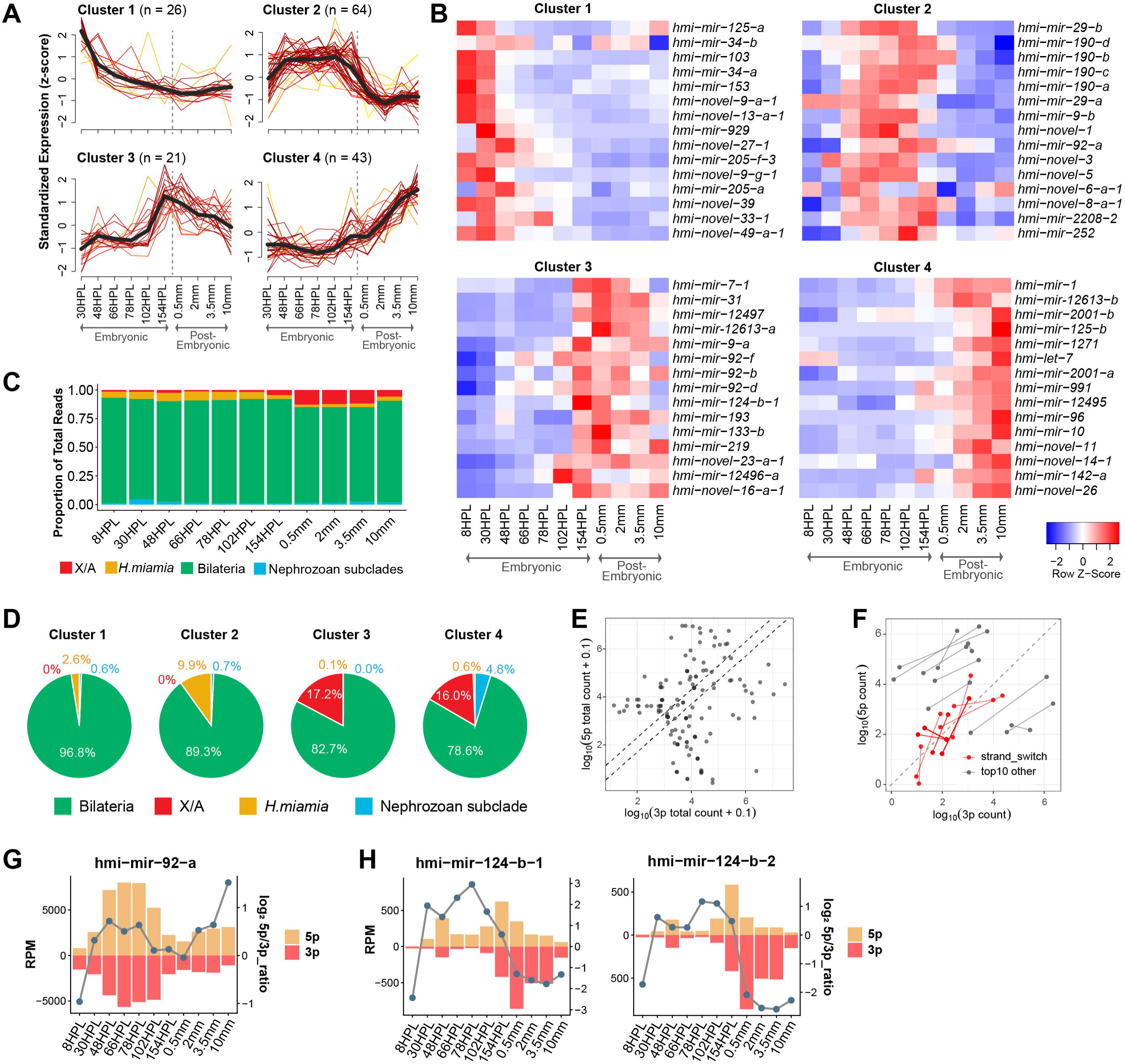
Developmental profiling of *H. miamia* miRNA expression. **A.** Fuzz c-mean clustering of miRNA expression across embryonic and post-embryonic developmental stages of the 154 highest-confidence miRNAs. Developmental stage information is provided in Figure 1C. The 8 HPL (early cleavage) stage was excluded in this analysis to enhance the clustering quality. The dashed line marks the transition between embryonic development and post-embryonic development. **B.** Heatmaps showing expression enrichment for the 15 most abundant miRNAs in each cluster. Within each cluster, miRNAs are ranked by total read counts. **C.** Proportions of miRNA reads in different evolutionary groups across development. The *H. miamia*-specific group is treated separately and is not included in the *Xenacoelomorpha* or the *Acoela* group. **D.** Proportions of miRNA reads assigned to each expression cluster in Figure 4A across different evolutionary groups. Percentages indicate the proportion of miRNA read counts from Bilateria-, Xenacoelomorpha/Acoela-, *H. miamia*-specific, and Nephrozoan subclade-derived miRNAs, in clockwise order. **E.** Scatter plot showing the log10 of total read counts from 3p (x-axis) and 5p (y-axis) strands of all miRNAs. Dashed lines indicate 3p/5p ratio of 2 and 0.5. **F.** Dumbbells plots showing developmental changes in strand preference. For each miRNA, the two points of dumbbell represent 3p and 5p read count coordinates at the two developmental stages showing the strongest 3p and 5p preferences, respectively. Stages with low expression (RPM <1) were excluded from this analysis. Red dumbbells indicate miRNAs with significant dominant strand alteration; note that some were overlapped on the plot. Grey dumbbells indicate the 10 most highly expressed miRNAs that did not exhibit dominant strand alteration. **G-H.** Expression profiles of 5p and 3p strands and their strand preference. Bars indicate normalized read counts (RPM; left y-axis) of the 5p/3p strands of *hmi-mir-92-a* (**G**) and *hmi-mir-124-b-1/2* (**H**), whereas points connected by lines indicate the 5p/3p ratio (right y-axis).

**Figure 5.**
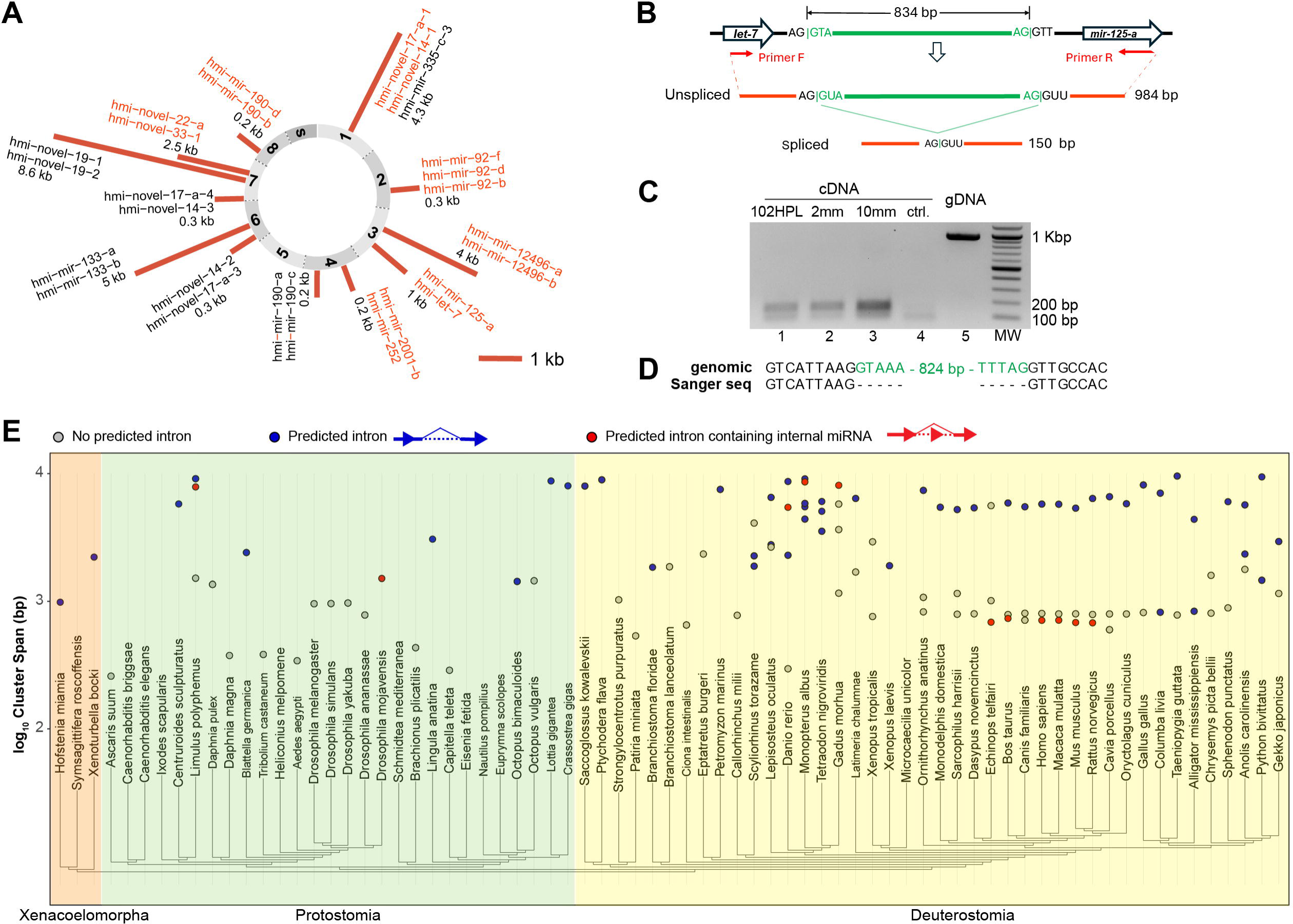
Splicing of the let-7 and mir-125-a primary transcript. **A.** Summary of the clustered highest-confidence miRNAs. Clusters are defined as groups of miRNAs spanning less than 10 kb. Bar lengths indicate the cluster length. Orange labels indicate the miRNAs in the clusters are likely co-transcribed based on the chromatin accessibility of putative promoter regions (Figure S11B-C). **B.** Genomic organization of the *hmi-let-7-hmi-mir-125-a* cluster and the proposed splicing of the primary miRNA transcript. **C.** RT-PCR products amplified from *H. miamia* cDNA (102 HPL/PL, 2mm/EJ, 10mm/SMA developmental stages, lanes 1-3), *C. elegans* mixed stage cDNA as control (lane 4), and *H. miamia* genomic DNA (gDNA, lane 5). **D.** Alignment of the predicted intron sequencing and Sanger sequencing results confirming the splicing event. **E.** Summary of the intron prediction of miRNA clusters consists of LET-7/MIR- 125/MIR-100 family members across bilaterian species. Y axis indicates log10 of the cluster span. Color-labeled points indicate clusters containing predicted introns with default ASSP thresholds. Red-labeled points indicate clusters in which the predicted intron contains one or more internal miRNA loci, such that splicing would remove these miRNAs from the primary transcript. Blue points indicate predicted introns without internal miRNA loci.

Among these clustered miRNAs, *the hmi-let-7* and *hmi-mir-125-a* are encoded within a 984 bp genomic region and appear to share a putative promoter, suggesting that they are co-transcribed as a single pri-miRNA transcript (Figure S11B). Interestingly, we detected a constitutive splice donor and a splice acceptor site between the two pre-miRNA hairpins, predicting the removal of an 834 bp intron from the primary transcript (Figure 5B). PCR amplification of the cDNA yielded an amplicon approximately 850 bp shorter than the amplicon from genomic DNA, consistent with the intron removal (Figure 5C). Sanger sequencing of the cDNA amplicon confirmed the predicted splicing event (Figure 5D). To test whether such intron removal is developmentally regulated, we chose three developmental stages: 102HPL stage, in which both miRNAs exhibited their lowest normalized expression; 10 mm stage, in which their expression is the highest; and 2 mm stage, at the onset of the peak expression. The spliced amplicon was detected at all three stages, indicating that this intron removal happens constitutively throughout tested *H. miamia* development (Figure 5C). Notably, we didn’t identify comparable splice donor and splice acceptor sites within the intervals between other predicted co-transcribed highest-confidence miRNA clusters. Together, these results indicate that *hmi-let- 7* and *hmi-mir-125-a* are co-transcribed and that their pri-miRNA transcript undergoes constitutive splicing.

The genomic clustering of members of the LET-7, MIR-100, and MIR-125 family has been reported in multiple species, including human, zebrafish, and *Drosophila* (49, 53). To investigate whether the pri-miRNA splicing we detected in *H. miamia* is also a conserved feature of these evolutionarily conserved miRNAs, we examined 73 bilaterian species containing members of these miRNA seed families. We identified 116 genomic clusters from 63 species in which two or more miRNAs with distinct seed sequences are located within 10 kb genomic intervals (Figure S12A). We then predicted the splice donor and splice acceptor sites within the intergenic regions between these clustered pre-miRNAs. We found that 63 clusters from 43 species, including Protostomia, Deuterostomia, and Xenacoelomorpha, contain donor and acceptor sites consistent with intron removal from the primary miRNA transcript (Figure 5E). Together, these results suggest that the pri-miRNA splicing within conserved LET-7/MIR-100/MIR-125 clusters is likely an evolutionarily widespread feature across Bilateria.

### *H. miamia* miRNA target sites prediction

To generate hypothesis for subsequent studies of miRNA functions in development and whole-body regeneration, we predicted the miRNA targets using the annotated 3’ UTR of protein-coding genes. Among the protein-coding genes, 89.7% contained annotated 3’ UTRs, with a mean length of 282 nt, consistent with the reported correlation between genome sizes and mean 3’ UTR lengths across animals (Figure 6A) (54). Note that for our subsequent analyses we only included 3’ UTRs with a length of more than 15 nt, the minimum length to incorporate miRNA binding (55, 56). We then predicted the target sites within these 3’ UTRs using our previously described algorithm, which identifies miRNA targets by evaluating both seed and 3’ non-seed pairing configurations (37, 57). In total, we predicted 97,127 target sites with perfect seed pairing (Table S6A) and demonstrated that 63.4% of all *H. miamia* protein-coding genes, and 71.9% of those with predicted 3’ UTRs, have the potential to be regulated by miRNA(s) (Figure 6B). The miRNA target sites are located in the 3’ UTRs with a mean distance of 179 nucleotides from the stop codons, displaying a distribution slightly closer to the stop codon compared to the overall predicted 3’ UTR lengths (Figure 6C). On average, a protein-coding gene is predicted to contain 7.7 miRNA sites (median 4), allowing it to be regulated by 5.1 miRNA seed families (median 3), while a single miRNA is predicted to be able to regulate 1729 targets (median 1363) (Figure 6D-E), which are comparable to other model species (Figure S13A-B). We also show that some evolutionarily conserved genes can be regulated by the same families of miRNA, despite variation of their 3’ UTR sequences. For example, we predicted five miRNA binding sites for four seed families of miRNAs in the 3’ UTR of lin28b, which encodes a putative *H. miamia* homolog of the evolutionarily conserved RNA-binding protein *LIN-28*. Among the four seed families, the sites for LET-7 family miRNAs and MIR-92 family miRNAs were also observed in other species, including humans and *C. elegans* (Figure 6F-G) (58).

**Figure 6.**
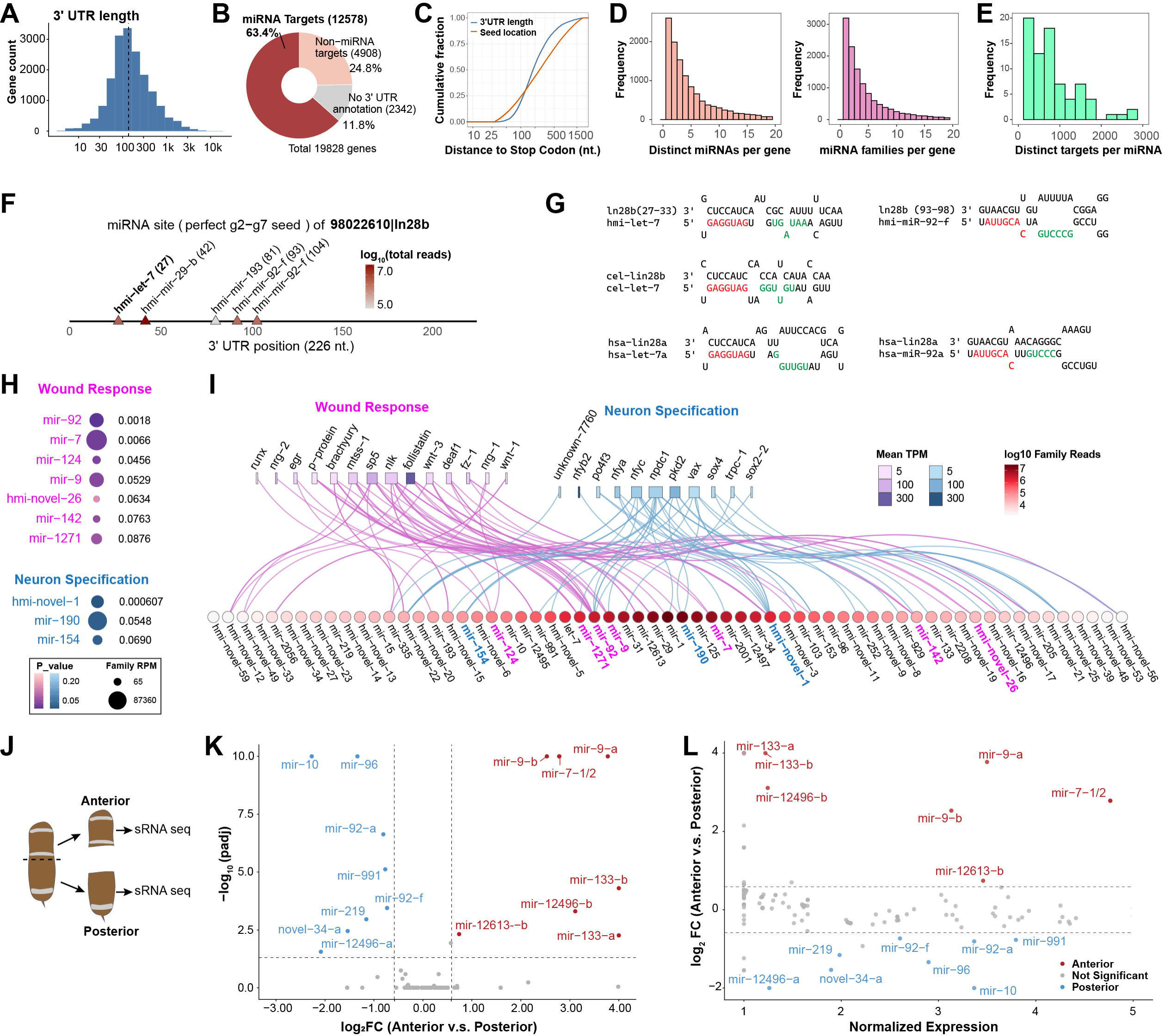
miRNA target prediction and hypothesis generation for *H. miamia* miRNA functions. **A.** Distribution of 3’ UTR length. Only the longest 3’ UTR was used for genes with multiple mRNAs. Dashed line, mean value. **B.** Proportions of *H. miamia* protein-coding genes categorized by predicted miRNA regulation status. Only miRNA target sites with perfect g2-g8 seed pairing are included in this analysis. **C.** Empirical cumulative distribution of the 3’ UTR lengths and miRNA seed binding site’s location (g2). **D-E.** Distribution of the number of predicted miRNA target sites and miRNA seed families per gene (D) and predicted targets per miRNA (E). Only target sites with perfect g2-g8 seed pairings were included in C-E. **F.** Location of predicted miRNA binding sites on the 3’ UTR of *H. miamia* homolog of *lin-28.* Color scale, total read counts of the miRNA family. Numbers, the location of the seed (g2). **G.** Base-pairing configurations of *let-7* and *lin-28*. Red letters, miRNA seed region (g2-g8). Green letters, critical non-seed region (g11-g16). **H.** Enrichment for target sites of each miRNA family in the wound response network (magenta) and the neuron specification network (blue). P-values, cumulative probability of one-tailed hypergeometric test. miRNA families that significantly enriched with p < 0.1 are shown. **I.** Putative miRNA-target interactome categorized by the wound response network and neuron specification network. Lines connecting miRNA seed families and protein-coding genes represent predicted target sites. Color scale, total read counts of the miRNA family. Bar lengths, number of distinct miRNA families predicted to regulate that gene. Bar color scale, mean TPM of the protein-coding genes calculated from previously published RNAseq datasets. **J.** Illustration of the *H. miamia* amputation strategy. **K-L.** Volcano plot and MA plot for differential expression analyses of all highest-confidence miRNAs between anterior and posterior fragments. Colored and labeled points, miRNAs that were differentially expressed with statistical significance (|FC| ≥ 1.5, p.adj ≤ 0.05).

It has also been shown that miRNAs can bind to target sites with imperfect seed paring and confer robust target repression, and such imperfect seed complementary requires compensatory pairing of the 3’ non-seed region (37, 38, 59–61). To this end, we identified 24,583 sites with imperfect seed pairing accompanied by 3’ pairing (Table S6B, Figure S13C-D). For this 3’ compensatory pairing configuration, the only target sites included were those with either a single GU base pair at g5-g8 or a single-nucleotide bulge on the target side of the seed helix, accompanied by at least three consecutive Watson-Crick pairing in the g11-g16 non-seed region (37, 38, 62, 63). We found 793 genes containing miRNA binding sites only with 3’ compensatory configuration, 5177 genes containing sites only with perfect seed pairing, and 7402 genes containing sites with both configurations (Figure S13E-H).

### Generating hypotheses for *H. miamia* miRNA targeting functions

To investigate the potential roles of *H. miamia* miRNAs in regulating biological processes, we analyzed their putative target interactions within two previously characterized gene groups: (a) the wound response gene set including the EGR-gene regulatory network and posterior patterning network and (b) the neuron specification gene set (18, 19, 46, 64). We focused on genes containing miRNA sites with perfect seed pairing as well as those with both perfect seed pairing and 3’ compensatory configuration (Figure 6H-I; Figure S13I-J). For the genes containing miRNA target sites with perfect seed pairing, we aggregated target sites by miRNA seed families because they can be regulated by any member of the family. For each miRNA seed family, we performed a hypergeometric test to assess target site enrichment in the two functional gene groups. Our results revealed significant enrichment (p<0.1) of the MIR-92, MIR- 7, MIR-124, MIR-142, MIR-1271 and HMI-NOVEL-26 families in the wound response group, while the HMI-NOVEL-1, MIR-190, and MIR-154 families were significantly enriched in the neuron specification gene group (Figure 6H-I).

Among the predicted miRNA targets in the wound response group included the genes *sp5, fz-1,* and *wnt-3* which have been previously described to exhibit asymmetric expression along the *H. miamia* anterior-posterior (AP) axis, with enrichment at the posterior of whole animals (13, 19). This prompted us to ask whether the correspondingly enriched targeting miRNAs are differentially expressed along the AP axis, and if so, whether their expression anti-correlates with that of the posterior genes. To address this, we performed sRNA-seq on the anterior and posterior fragments following transverse amputation (Figure 6J, Table S7). We found that both miRNAs in the *H. miamia* MIR-9 family, which are predicted to target the posterior-expressed genes *sp5* and *wnt-3*, exhibited significantly enriched expression in the anterior fragment (Figure 6K-L). Note that the anterior-expression pattern of MIR-9 miRNAs has also been observed in other animal species, including mice, frogs, and zebrafish (65–67). Additionally, hmi- miR-7-1/2, one of the most abundant *H. miamia* miRNA family and a putative regulator of posterior-expressed *fz-1* was also highly enriched in the anterior fragment (Figure 6K-L). The anti-correlated expression along the AP axis of MIR-9 and MIR-7 miRNAs with *sp5*, *fz-1*, and *wnt-3*, suggests that these miRNAs can be candidate genes for future study as potential regulators of posterior patterning in *H. miamia* development and regeneration.

## DISCUSSION

Our comprehensive annotation of miRNAs in *Hofstenia miamia* expands the current understanding of xenacoelomorph miRNAs through three key advances: substantially deeper small RNA sequencing than previous studies, comprehensive characterization of miRNA expression dynamics across developmental stages, and a more complete genome assembly with rigorous scoring of predicted miRNA quality. The increased sequencing depth improves the completeness of miRNA annotation because the *de novo* miRNA prediction algorithms used in our study, sRNAbench and miRDeep2, require sequencing reads mapping to both the mature and star strands of a candidate hairpin to identify a pre-miRNA. Consequently, miRNAs with low overall expression or strong strand bias may escape detection if the star strand is insufficiently represented. Consistent with this limitation, among species with extensive small RNA sequencing resources, the number of annotated miRNA loci exhibits a positive correlation with genome size, whereas species with limited sRNA sequencing resources frequently fall below this trend, suggesting incomplete miRNA annotation in those cases (Figure S6B). Beyond sequencing depth, the inclusion of a comprehensive range of developmental stages is also critical for robust miRNA annotation. For example, our analysis identified many miRNAs enriched at early embryonic stages (e.g., Hmi-miR-34-a) or late post-embryonic stages (e.g., Hmi-miR-2001-b) (Figure 4B). Although these two miRNAs were highly expressed at their respective stages and exhibited high-confidence structural features, they were not detected in the previously published annotations, likely due to the lack of samples from those developmental stages. Lastly, the improved genome assembly contains substantially fewer assembly gaps, reducing the likelihood that miRNA loci are fragmented, collapsed, or missing from the reference genome and thereby improving the completeness of miRNA annotation.

False-negative miRNA annotations discussed above can have consequences beyond simply underestimating miRNA repertoire size; they can also bias comparative and evolutionary analyses. For example, in Figure 4C-D we showed that the xenacoelomorph/acoel-specific miRNAs appear to be predominantly expressed during post-embryonic development, which may suggest that these miRNAs have primarily post-embryonic functions. However, an alternative explanation is that some miRNAs that are currently categorized as *H. miamia*-specific with expression enrichment during embryogenesis, are actually conserved in *X. bocki* or *S. roscoffensis.* They may simply have escaped annotation because the available datasets for these species lacked sufficient sequencing depth and/or comprehensive embryonic sampling. In this scenario, these miRNAs would be incorrectly classified as *H. miamia-*specific because of false-negative annotations in *X. bocki* and *S. roscoffensis*, thereby biasing the inferred relationship between miRNA evolutionary origin and developmental expression. This example illustrates the importance of comprehensive miRNA annotation for accurate comparative, evolutionary and functional analysis.

*De novo* miRNA annotation can also generate substantial numbers of false positive predictions. Among the 545 candidate miRNA loci from our study, some may represent non-canonical small RNAs. For example, several loci exhibited sub-optimal predicted hairpin structures, suggesting biogenesis mechanisms that do not rely on a stable hairpin structure characteristic of *bona fide* miRNAs. One such example is the abundantly expressed Hmi-miR-205-e-1 (36,530 total reads; top 12% by abundance). Its predicted structure displayed minimum free energy (MFE) of -13.2 kcal, suggesting a hairpin structure with relatively low stability (68). Meanwhile, many candidate loci were supported by only a small number of reads, raising the possibility that they represent degradation products of other classes of stem-loop-derived small RNAs rather than *bona fide* miRNAs. By convention, such low-confidence predictions are often excluded from being reported or curated. Nevertheless, we reasoned that excluding genomically encoded small RNA-producing loci solely on basis of arbitrary structural or expression thresholds could diminish the value of this resource for future studies. Therefore, we chose to report all 545 predicted miRNA loci together with their structural and expression properties in the annotation GFF file. This approach allows researchers to access the complete small RNA dataset and utilize the information as they see fit for their specific analytical needs.

To facilitate the use of the *H. miamia* miRNA annotation in both our downstream analyses and for future studies, we developed a scoring system that provides three complementary levels of information. First, we report the raw values of all scoring features, including structural features, expression levels, and prediction quality metrics, which allow users to independently evaluate each candidate locus. Second, we assign penalties to individual features based on comparisons with curated bilaterian miRNAs or biologically motivated thresholds, thereby providing a standardized measure of annotation confidence for each feature. Third, we integrate these evaluation into a final annotation score that summarizes the overall confidence of each predicted miRNA locus. In parallel to employing the scoring system to assign annotation scores to the set of 545 candidate miRNA loci, we also defined a highest-confidence subset of 154 miRNA loci. The parameters and thresholds used to generate this highest-confidence subset were determined using curated miRNAs as a reference, applying stringent filtering criteria, and were designed to align with the current curation principles of MirGeneDB (Table S2; Materials and Methods). Together, the annotation score and the highest-confidence subset provide complementary resources, enabling users to apply filtering criteria appropriate for their specific application while maintaining complete access to the underlying annotation data. Rather than applying a single hard threshold to define *bona fide* miRNAs, our framework preserves the underlying annotation evidence and allows users to apply confidence thresholds appropriate for their own biological questions. We anticipate that this framework will be broadly useful for *de novo* miRNA annotation in other non-conventional model organisms.

Notably, although miRNAs excluded from the highest-confidence subset generally exhibited lower scores and expression abundance (Figure S6B), several of these loci nevertheless performed relatively well across most evaluation criteria and were excluded from the highest-confidence subset because they failed a single stringent criterion. This indicates that the highest-confidence subset should not be interpreted as a definitive boundary between *bona fide* and non-*bona fide* miRNAs. For example, *hmi-mir-154-a* has 424,829 total reads (top 0.6% by abundance) and an annotation score of 65, but was excluded because its 3’ overhang (4 nt) exceeded the defined threshold (3 nt). Similarly, Hmi-miR-92-c and Hmi-miR-92-e, both belonging to the conserved bilaterian MIR-92 family, were excluded because of their relatively low mature-strand base-pairing ratio (0.59), despite their high expression levels (939,836 and 439,772 total reads, respectively). In addition, *hmi-mir-92-c* is clustered with three other MIR-92 family members that were included in the highest-confidence list, further supporting its authenticity. For this study, we intentionally avoided adjusting thresholds or manual manipulation on a locus-by-locus basis for the highest-confidence subset to be objective and stringent.

Nevertheless, we anticipate that some currently excluded loci may be promoted to the highest-confidence category as additional datasets become available, such as tissue-specific small RNA sequencing, comparative genomics, Argonaute pull down, or evolutionary conservation analysis of target sites.

Our data provide evidence supporting certain revisions to the current assignment of the phylogenetic origins of several miRNA families. In the current version of MirGeneDB (v3.0), MIR-103 was assigned to the Bilateria node, whereas nine families (MIR-8, MIR-71, MIR-76, MIR-184, MIR-216, MIR-242, MIR-281, MIR-315, MIR-375) were reassigned from Bilateria to Nephrozoa based on the absence of these families in the available xenacoelomorph small RNA datasets (Figure S7F). Our expanded annotation fully supports these revisions, as we detected the MIR-103 family in *H. miamia* but did not detect any members of the above nine families reassigned to *Nephrozoa-*specific. In contrast, 38 of our highest-confidence miRNAs (comprising 10 seed families) in our *H. miamia* datasets, are currently annotated as specific to nephrozoan subclades in MirGeneDB (v3.0), and seven of them are present in multiple non-sibling species (Figure 3E). These seven seed families (MIR-15, MIR-205, MIR-142, MIR-335, MIR-154, MIR-929, MIR-1271) may warrant being reassigned to the Bilateria node, which would increase the number of Bilateria node miRNAs to 33. Most of these families contain miRNAs with high annotation scores and moderate expression levels (Figure 3F). For example, in our annotation, the MIR-205 family contains 15 annotated loci, and three family members are included in the highest-confidence subset with a total of 19,543 reads and the highest annotation score (100/100) for all three miRNAs. Notably, our developmental profiling shows that the MIR-205 family is expressed almost exclusively during early embryonic stages (Figure 4B). This restricted expression pattern likely explains why MIR-205 escaped previous *H. miamia* annotations, which lacked comprehensive sampling of early embryonic samples. These findings demonstrate that phylogenetic assignment of miRNA families remains sensitive to annotation completeness, particularly in early diverging animal lineages where in-depth and comprehensive developmental small RNA sequencing datasets remain limited.

Among ancestral bilaterian miRNAs, the MIR-1 and LET-7 families both contain one family member that shows the highest level of phylogenetic sequence conservation in the 3’ non-seed region. In *H. miamia,* we show that Hmi-miR-1 maintains the same level of non-seed conservation as other bilaterian MIR-1 miRNAs. In contrast, *H. miamia* does not contain the deeply conserved *let-7a* miRNA and instead possesses one *let-7* family miRNA which differs from *let-7a* at g10-g18 (Figure 3I). Interestingly, the 3’ distal sequence (g19-g22) of this *Hmi-let- 7* is nevertheless identical (AGUU) to the deeply conserved *let-7a* (Figure 3I). The conservation at the g19-g22 region was also observed for some LET-7 family miRNAs that are not conserved with *let-7a* at g10-g18, for example, *Sme-let-7d* (Figure 3I). Given that pairing at the 3’ distal region is critical for target-directed miRNA degradation (TDMD), we propose that the sequence conservation of g19-g22 at *Hmi-let-7* might be driven by evolutionarily conserved mechanisms regulating *let-7* family miRNA stability (39, 69).

We observed that thirteen highest-confidence miRNA genes expressed both the 5p and 3p strands at similar levels across more than half of the sampled developmental stages, suggesting that both strands may be functional and regulate distinct target sets, for example *hmi-mir-92-a* (Figure 4K). In addition, several miRNAs exhibited developmentally regulated switches in strand preferences, for example *hmi-mir-124-b-1* and *hmi-mir-124-b-2* (Figure 4J, Figure S9A). Such developmental changes in 5p/3p strand preference may reflect multiple underlying mechanisms: First, different tissues or cell types may preferentially express either the 5p or 3p strand, so that the enrichment or emergence of such tissues or cells may lead to the changes in the overall 5p/3p ratio. Second, developmental regulation within the same cell type may influence the strand selection machinery, resulting in temporal changes in strand preference. Third, changes in strand preference may arise from differential expressions of miRNA loci that produce identical mature sequences from one arm but distinct sequences from the other. This scenario is illustrated by the MIR-124 family, in which the 3p strands of *hmi-mir-124-b1* and *hmi-mir-124-b-2* are identical and evolutionarily conserved, whereas the 5p strands have different sequences and show little conservation (Figure 4H). Consequently, an increase of the *mir-124-b-1-3p* strand could either come from its own locus, or from the *mir-124-b-2* locus with the increased expression of *mir-124-b-2-5p* to regulate its own targets. Currently, we do not have any means to distinguish between these and other hypothetical mechanisms. For these reasons, we recommend exercising caution when interpreting changes in 5p/3p strand preferences.

In *H. miamia,* we found that an intron is spliced from the transcript containing *hmi-let-7* and *hmi- mir-125-a*, and our comparative analysis suggests that splicing within clustered LET-7/MIR- 125/MIR-100 family members may be widespread across Bilateria. The predicted splice sites appear to be robust, as 86% of the predicted introns were retained even when the splice donor and acceptor score thresholds were increased to the highest values available (Figure S12B) (70). Although the molecular and biological significance of this splicing remains unclear, our analysis suggests several potential hypotheses. One possibility is that the splicing allows the expression of individual miRNAs within a cluster to be uncoupled. In 11 species, LET-7/MIR- 125/MIR-100 family clusters contain more than two miRNAs with at least one miRNA located within the predicted intron (Figure 5E). In these clusters, splicing would therefore remove the internal miRNA from the transcript, potentially allowing its expression to be uncoupled from the distal miRNAs. This mechanism raises the possibility that splicing provides an additional regulatory layer within the clustered LET-7/MIR-125/MIR-100 family members. Moreover, the splicing substantially shortens the primary miRNA transcripts, which may influence the subsequent processing of the two miRNAs. In *H. miamia*, *hmi-let-7* and *hmi-mir-125-a* exhibited strongly correlated expression both temporally across development (Figure S12C) and spatially between the head and tail fragments (Figure S12D). Despite this coordinated expression pattern, the stoichiometry of the two mature miRNAs was consistently imbalanced, with *let-7* approximately sixfold less abundant than *miR-125-a*. One possibility is that the intron removal alters the spacing and structural context of the two pre-miRNA hairpins, thereby favoring Microprocessor binding or cleavage at one hairpin over the other. Cryo-EM structural analysis of human Drosha-DGCR8-RNA indicates that the Microprocessor occupies a substantial volume around its bound hairpin substrate (71). Following the splicing, the let-7 and mir-125-a hairpins are separated by approximately 54 nt between the predicted DGCR8-binding regions. Although a fully extended RNA would theoretically provide sufficient space to accommodate Microprocessors complexes at both hairpins, RNA secondary-structure prediction indicates that the intervening sequence has the potential to form secondary structures that could reduce the spatial separation between the two pre-miRNA hairpins and potentially disfavor simultaneous processing of both hairpins (Figure S12E). Whether and how this transcript architecture contributes to the unequal accumulation of mature *let-7* and *miR-125-a* remains unknown.

Genome editing of the splice donor and acceptor sites will be required to determine whether splicing directly influences the processing or regulation of the clustered miRNAs in both *H. miamia* and other species.

To generate hypotheses for the *in vivo* functions of *H. miamia* miRNAs, we suggest that criteria for prioritizing miRNAs and target genes for functional studies could include the following: (1) miRNAs that are enriched for cognate target sites within the gene cohort associated with the biological functions of interest; (2) miRNAs and predicted targets that display spatially distinct and/or developmentally dynamic expression patterns consistent with the developmental process of interest; (3) miRNAs and targets with anti-correlated expression patterns. For example, in our study, we found that predicted targets of the MIR-9 family were enriched in the gene group associated with posterior patterning, including *wnt-3*. Moreover, the MIR-9 family miRNAs were preferentially expressed in the anterior fragment of the animal and anti-correlated with *wnt-3*, which is expressed in the posterior and regulates the AP axis re-patterning in regeneration according to previous studies (19). These findings suggest that MIR-9 miRNAs are compelling candidates for the study of the roles of miRNAs in AP axis re-patterning during regeneration. It is noteworthy, however, that a single miRNA can contribute to multiple biological functions by repressing multiple targets, which can confound the interpretation of miRNA gene loss-of-function phenotypes. This concern can be addressed by generating mutants with miRNA binding sites disabled from the 3’ UTR of candidate target genes to directly test the functional contributions of candidate miRNA-target interactions.

For subsequent functional studies of the annotation miRNAs, we conducted target site prediction for *H. miamia* miRNAs. While these predicted miRNA-target interactions may contain false positives, given that the miRNA-mediated repression is influenced by numerous factors, our target predictions nevertheless provide a catalog of hypothetical gene regulatory interactions of the *H. miamia* miRNAs (1, 4, 72) and hence establish a foundation for future functional studies of the *H. miamia* miRNAs. Given that CRISPR genome editing has been successfully demonstrated for *H. miamia* (17), we anticipate that the biological roles of the *H. miamia* miRNAs and their predicted target interactions can be investigated by generating miRNA and target mutants and examining their phenotypic effects.

## CONCLUSION

In this research, we established comprehensive genomic and miRNA resources for *H. miamia,* including a PacBio-based genome assembly, updated protein-coding gene annotation, and the annotation of 545 miRNA loci, including a subset of 154 highest-confidence miRNAs supported by extensive developmental small RNA sequencing. We further developed an objective scoring framework for evaluating miRNA annotation confidence, providing a strategy that may facilitate miRNA annotation in other emerging model organisms. Using these resources, we characterized the evolutionary origins and developmental expression dynamics of the *H. miamia* miRNA repertoire, identified splicing associated with clustered LET-7/MIR-125 miRNAs, and generated a genome-wide map of putative miRNA-target interactions. Together, these resources and analyses establish a foundation of investigating miRNA-mediated post-transcriptional regulation during *H. miamia* development and whole-body regeneration, and provide a framework for comparative studies of miRNA evolution and functionality across animals.

## METHODS

### Animal culturing and sample preparation

*H. miamia* animals in this study were taken from a colony derived from an initial population collected in Bermuda (13). Embryos and hatched animals used here were the progeny of random mating. Embryos and hatched animals were cultured in conditions previously described (13, 73). Early cleavage embryos were defined as embryos from zygote to 8-cell stages. Samples of all other embryonic stages were collected from synchronized early cleavage embryos, and the development stages were determined by the time post egg-laying as described in Figure 1B, and the developmental stages were verified by embryo morphology described in (13, 73). The hatchling juveniles were collected from individual animals newly hatched within 14 hours. Young juvenile, late juvenile, and sexually mature adults were categorized based on their size from head to tail as described in Figure 1B, and samples were collected after 6 days of starvation. The sexually mature adults were further visually confirmed to contain oocytes. The late juvenile-sized animals were used for the anterior/posterior fragments after transverse amputation. Harvested embryos or worms were collected in 1.7 mL Eppendorf tubes, most of the artificial seawater (ASW) was removed by pipetting, and samples were flash-frozen by incubating on dry ice for 15 min and stored at -80 °C.

### PacBio genome assembly - Genomic DNA preparation and sequencing

The genomic DNA (gDNA) for the PacBio genome assembly was isolated from the same Head3 animal from which the previous genome was assembled. The P0 Head3 animal was starved for 10 days to minimize contamination from residual food DNA. The animal was then amputated and gDNA was extracted from the whole tail fragment using NEB Monarch Spin gDNA Extraction Kit (NEB Cat# T3010S). gDNA was eluted by 85 µl elution buffer (10 mM Tris-HCl, pH 9.0, 0.1 mM EDTA). The gDNA concentration was measured using Qubit dsDNA HS kit (ThermoFisher Cat# Q32851) to be 33.8 ng/µl. The gDNA quality was assessed by Nanodrop (OD_260/280_ = 1.85). gDNA was further purified using Beckman Coulter AMPure beads, integrity was confirmed by TapeStation, with peaks observed at 59 kB. The PacBio library preparation and sequencing were performed at Harvard Baurer Core Facility using PacBio Revio.

### PacBio genome assembly - Genome assembly and scaffolding

For initial assembly, 28.2 Gb of high quality PacBio Hi-fidelity reads were generated for an estimated 30X coverage and assembled using hifiasm v0.19 (74, 75). To increase contiguity, HiC reads were mapped against the draft assembly using bwa mem, and custom scripts were used with samtools to filter and combine mapped reads, then the draft assembly was scaffolded using YaHS v1.2 (76). RagTag was used to scaffold against a previously existing assembly generated from Nanopore data to further increase contiguity, verifying the joints made by remapping the Nanopore data using minimap2 (77, 78). Subsequently, the external contaminants were determined by using both NCBI Foreign Contaminant Screen tool and by aligning the *H. miamia* preliminary assembly to all currently curated genomes (32,039 genome and 11.4M scaffolds/chromosomes) (79–81). 11 scaffolds were identified as external contaminants, which were either manually removed or masked in the assembly before doing one final round of scaffolding (Figure S1B). With the .hic contact map file generated from YaHS, the Juicer HiC toolkit were used in combination with JBAT Juicebox visualizer to manually curate the assembly, identifying and correcting several mis-joins and inversions in the 8 chromosomal scaffolds (primarily in centromeric and telomeric regions) (82, 83). The final version of the assembly is highly contiguous, with a scaffold N50 > 130Mb and 90% of the assembly sequence contained in the 8 chromosome scaffolds. Genome completeness was measured using compleasm with the ‘metazoa_odb12’ and ‘eukaryota_odb12’ core gene datasets, which resulted in 80% (75% single, 1.6% duplicated, 3.6% fragmented) and 91% completeness (88% single, 0% duplicated, 3.1% fragmented), respectively (84). The final assembly and the previously published short-read assembly were then subjected to QUAST for statistical analysis (85).

### Genome annotation - StringTie annotation

The PacBio genome was annotated using 651 M paired-end non-directional RNAseq reads from 14 experimental conditions spanning *H. miamia* embryonic and post-embryonic stages. The PacBio genome assembly was first indexed *by STAR v2.7.10a with*out splice-junction (SJ) files. RNAseq reads were mapped, yielding an 85.43% overall mapping rate and generating SJ information. The PacBio genome was then re-indexed using the SJ files, and the same RNAseq reads were mapped again with *--outSAMstrandField intronMotif* to include the *-XS* tag in the output. This secondary mapping resulted in 14 BAM files (one per experimental condition) with a total mapping rate of 85.14%. Each BAM files was subjected to *StringTie/2.2.1* with option *-s 3.75* to generate a GTF file, and all 14 GTF files were then merged into a final StringTie annotation file by *StringTie –merge* with default parameters, yielding in 27,756 gene loci and 57,042 mRNAs. TPM and per-Base coverage were calculated by *StringTie -e -B* (21, 86).

### Genome annotation - Recovery and integration of missing genes using reference transcriptome

We observed that some genes present in our previously published transcriptome were absent in the StringTie annotation, including genes that have been functionally characterized. To address this, we integrated gene models from our latest reference transcriptome (22). Specifically, a total of 27,805 transcripts from the reference transcriptome were mapped to the PacBio genome assembly by *minimap2*, with a GTF file using SAM2GTF (77). Of these, 27,416 transcripts (98.6%) mapped to the PacBio genome, and 26,588 transcripts (95.6%) were confidently propagated as gene loci. The resulting GTF files from StringTie and the processed reference transcriptome were compared using GFFcompare, with the StringTie annotation as the reference (87). Genes from the reference transcriptome assigned to matching classes “i", “y”, “p”, “o”, and “u” were retained and filtered for TPM > 0.5 (top 95th percentile of all genes). 636 class “o” genes were further removed if a higher-class code was assigned to another reference gene. Remaining genes from the reference transcriptome were incorporated into the StringTie annotation, resulting in 29,836 genes in the final genome annotation. One gene, 98200327|at2b3-3 was manually split into 98200327|at2b3-3 and 98041185|po2f3 based on our unpublished functional study.

### Genome annotation - ORF and UTR annotation

To predict ORFs, a transcriptome FASTA file was generated from the final annotation by *TransDecoder/util/gtf_genome_to_cdna_fasta.pl* . The output GTF was converted to GFF3 by *TransDecoder/util/gtf_to_alignment_gff3.pl*, and a gene_to_transcript roadmap file was generated by a custom script. The longest ORF per locus was predicted by *TransDeocder.LongOrfs*, and the output were subjected to homology searches via BLASTP against human proteome (Uniprot *UP000005640*) with a cutoff of 1e-3. The best ORF for each gene locus was selected by *TransDecoder/TranDecoder.Predict.* Predicted ORFs were then propagated to the genome by *Transdecoder/cdan_alignment_orf_to_genome_orf.pl* with additional custom scripts for naming suffix. UTR features < 3 bp in length and associated with incomplete ORFs were removed (exon features unchanged). The final GFF3 was converted to GTF by *AGAT*, with a “biotype=protein_coding” flag added to each gene with predicted ORF. Transcripts without ORF predictions (12,877 transcripts) and those with inconsistent strand propagation were manually added and annotated as “biotype=non-coding” and “biotype=protein_coding_antisense”, respectively (86). The mRNA sequences were then exported and subjected to BUSCO analysis.

### Genome annotation - Strand assignment and final gene identification

The annotation included 5,505 single-exon non-coding genes without stranding assignment. To determine strand information, 607 million unpublished directional single-ended RNAseq reads were mapped to the PacBio genome using STAR, and gene counts were quantified for both strands using subread/featureCounts. Strand bias was evaluated as log_2_(plus_count + 0.1)/(minus_count + 0.1)), and a strand bias cutoff was determined by performing a split test with the genes that already had strand annotation. Strand information was assigned to previously unstranded genes if the dominant strand bias exceeded 2.95 as determined by the split test and had more than 50 reads. This approach resulted in strand assignment for 1,573 genes.

Gene IDs and gene names in the new annotation were then assigned by running GFFcompare with the new annotation as subject and the reference transcriptome as the reference. Genes in classes “=”, “j”, “k”, “c”, “m”, “n” inherited the gene_id and gene_name from the reference transcriptome directly, with overlapping genes de-duplicated using custom script. Remaining genes were assigned with gene_id prefixed by 983XXXXX|, and gene names assigned based on the homology search results. Genes without human homologs (identified via ORF BLAST) were designated as "unknown-" in their gene names. For genes carried over from the reference transcriptome, the original "unknown-" suffix was retained to ensure unique nomenclature.

Finally, the gene_id and gene_name of 135 published and functionally characterized genes were manually curated based on previous publication.

### Total RNA preparation

The animal samples were thawed and lysed by adding 600 µL QIAzol (Qiagen, Cat: 79305) and shaking on vortex at room temperature for 15 min. Total RNA was extracted by adding 200 µL ddH_2_O, 175 µL chloroform, vortex at room temperature for 1 min, centrifugation at 25,000 rcf for 10 min at 4 °C, and recovery of the aqueous phase. The aqueous phase was then re-extracted with 1X volume phenol:chloroform:isoamyl alcohol (25:24:1, pH = 6.3), followed by a second re-extraction with 200 µL chloroform to remove residual phenol. Total RNA was then precipitated by adding 0.03 volume of 3M NaAc (pH = 5.5), 1.2 volume of isopropanol, and 1.5 µL GlycoBlue (Invitrogen, Cat: AM9516), followed by incubation at -80 °C for at least 30 min, and recovery by centrifugation at 25,000 rcf for 30 min at 4 °C. Supernatant were discarded, and RNA pellets were subsequently washed twice with 75% ethanol, air-dried at room temperature for 5 min, and dissolved in RNase-free water.

### Small RNA sequencing – Library preparation

RNA samples were mixed at 1:1 (v/v) ratio with 2X RNA loading dye (95% (v/v) formamide, 0.01% (m/v) SDS, 0.05% (m/v) bromophenol blue, 0.05% (m/v) xylene cyanol, 0.5 mM EDTA), denatured at 85 °C for 90 sec, chilled on ice, and separated by 15% PAGE using 1X TBE buffer with microRNA marker (NEB, Cat: N2102S) as size maker. The gel was stained by 1:10000 (v/v) Sybr Gold (Invitrogen, Cat: S11494) in 1X TBE buffer with low-speed orbital shaking at room temperature for 8 min, and gel slice containing small RNA with approximately 18-26 nt was excised according to the size marker, ground by an RNase-free pellet pestle (Fisher Scientific, Cat: 12-141-364), and flash-frozen by dry ice for 30 min. Small RNA was extracted by adding 500 µL of 300 mM NaAc (pH = 5.5), 1 mM EDTA, and 0.25% (m/v) SDS, mildly shaking overnight at room temperature (Ingolia et al., 2012). The gel granules were excluded using a Spin-X 0.2 µm tube filter (Millipore, Cat: CLS8160), and the small RNA was concentrated by ethanol precipitation as described above, dissolved in water, and stored at -80 °C. For developmental samples, sRNA seq libraries were prepared using QIAseq miRNA Library Kit (Qiagen, Cat: 331565) with miRNA Index Kit Version 1 (Qiagen, Cat: 331592) following the manufacturer’s instructions except that the amplifying PCR was conducted with 10-12 cycles, and libraries were quantified by KAPA Library Quantification Kit (Roche, Cat: KK4824) and Qubit dsDNA HS Reagent (Invitrogen, Cat: Q32851) and sequenced using Illumina NextSeq 550 platform. For anterior/posterior samples, sRNA seq libraries were prepared and sequenced identically to the development samples, except for using the miRNA Index Kit UDI Version2 (Qiagen, Cat:331905) for barcoding and the NextSeq 2000 system for sequencing.

### Small RNA sequencing –Data processing

Small RNA reads were checked for quality before and after filtering using FastQC v0.11.9 (https://www.bioinformatics.babraham.ac.uk/projects/fastqc). The raw reads were deduplicated based on the 12-digit unique multiplexing unit (UMI) using UMI-tools v0.3.4 (88). The adaptor sequences were trimmed from the 3’ end of the raw reads, and the trimmed reads were quality-filtered with *-q 20* option and size-filtered with *-m 16 -M 28* options by Cutadapt v4.1 (89). The contaminating rRNA and tRNA reads were removed by mapping the filtered reads to indexed *H. miamia* rRNA and tRNA sequences curated in RNAcentral v22 (90) using Bowtie2 v2.5.0 (91, 92).

### miRNA annotation – primary miRNA prediction

For miRNA prediction, filtered reads from each library were independently processed using sRNAbench and miRDeep2 with default parameters, using the PacBio *H. miamia* genome as reference (26, 27). miR-451-like predictions were discarded. Predictions were further tracked across developmental stages: miRNAs predicted at only one developmental stage were retained only if detected in at least two of the three biological replicates at that stage. The remaining predictions from sRNAbench and miRDeep2 were then merged using BEDTools v2.30.0 (93), with each retained locus annotated according to which tool(s) supported its prediction. The pre-miRNAs were then subjected to RNAfold in Viennarna v2.6.3 to predict the secondary structure and minimum free energy (MFE) (94, 95). The max bulge asymmetry, mature base-paring ratio, 3’ overhang, and apical loop sizes were calculated from the bracket- dot structures from the RNAfold output.

### miRNA annotation – miRNA expression and in-cluster ratio evaluation

The genomic coordinates of the primary prediction of 690 loci were exported, and expression of both 5p and 3p strands were re-evaluated. To do this, the filtered reads from each library were mapped to the re-indexed PacBio genome by star v2.7.10a using default parameters (96). Read counting was done by featureCounts from subread v1.6.2 with reads mapped to multiple miRNA loci counted with *-M --fraction* options (*97*). The dominant strand (mature strand) is then defined as the strand with the highest global read abundance.

To calculate the in-cluster ratio, each pre-miRNA locus was expanded by 10 bp upstream and downstream, and the total number of reads mapping within the expanded region was quantified as described above. The in-cluster ratio was calculated from the mature-strand read count (Figure S3A) as defined previously (28) or from the combined 3p- and 5p-strand read counts (Figure 2H), divided by the total read counts within the expanded pre-miRNA locus after adding a pseudocount of 0.1. The Figure 2H formulation of in-cluster ratio is adopted to accommodate potential strand-switching events, where both 5p and 3p strands may be functional.

### miRNA annotation – False-positive filtering

To remove false-positive predictions, predicted miRNA loci were sequentially screened using the following criteria: (1) the pre-miRNA sequence could not be folded into a secondary structure (17 loci removed); (2) the mature-strand base-pairing ratio was below 0.5 (16 loci removed); (3) the predicted secondary structure did not represent a canonical pre-miRNA hairpin, for example forming a multi-armed hairpin structure (8 loci removed; Figure S2D); (4) the predicted locus represented an artificial assembly of the mature and star strands from two nearby pre-miRNA loci (1 locus removed; Figure S2E); (5) the combined mature- and star-strand in-cluster ratio was below 50% (12 loci removed); (6) the total fractional read count of either the mature or star strand was below 1 following read count re-evaluation (83 loci removed); (7) no individual library contained more than one mature-strand read after the fractional read count re-evaluation (8 loci removed). The numbers of removed loci at each step were counted sequentially; therefore, loci excluded by an earlier criterion were not included in the counts for subsequent criteria.

### 5’ heterogeneity analysis

The 5’ heterogeneity of all 545 annotated miRNAs was quantified using a custom R script. Briefly, the annotated 5’ end of each mature miRNA (i.e., as annotated by sRNAbench and miRDeep2) was used as the reference coordinate, and the genomic 5’ end of each aligned read was compared with the original reference coordinate to determine the 5’ shift. Reads with shifts between -3 and +3 nucleotides were included in the analysis. For each mature miRNA, 5’ heterogeneity was calculated as the percentage of reads with non-zero 5’ shifts relative to the total reads mapped to the locus.

High-abundant 5’ isomiRs were subsequently identified for reporting and for revising 5’ sequence annotation, as appropriate based on their global or developmental abundance. An isomiR was retained due to global abundance if it satisfied the following criteria: (1) global relative abundance ≥10%; (2) at least 100 global reads combining all libraries; and (3) at least 20 reads in at least one library. This identified 144 isomiRs from 124 miRNAs. To identify developmentally enriched isomiRs, an isomiR was retained if it satisfied all of the following criteria: (1) relative abundance ≥10% at a single developmental stage in at least two of the three biological replicates; (2) at least 100 total reads across all libraries; and (3) at least 20 reads in at least one library. This identified 136 isomiRs from 118 miRNAs. In total, 158 unique isomiRs from 136 miRNAs were identified (Table S4).

### Sequence revision of 5’ heterogeneous miRNA

The annotated 5’ sequence of the mature strand (reference miRNA, ref-miR) was revised when a 5’ isomiR satisfied any of following criteria: (A) The global abundance of the 5’ isomiR was greater than 50% of the total read count of the locus, or was greater than the total read count of the ref-miR (FC iso/ref > 1). (B) The isomiR had significantly higher expression (FC >2, p <0.05 in t-test) than the ref-miR at the developmental stage with the highest expression of the miRNA. (C) The isomiR had significantly higher expression (FC > 1.5, p < 0.1 in t-test) than the ref-miR in at least total 6 of the 11 developmental stages, or in at least three consecutive stages. (D) The 5’ end if the 7mer seed (g2-g8) of the isomiR corresponded to a seed family curated as an earlier node of origin, or to an identified highest-confidence *H. miamia* miRNA seed family (Figure S6A).

A total of 31 isomiRs from 25 miRNA loci satisfied the above criteria, including four miRNAs with multiple annotated 5’ isomiRs. In these cases, the isomiR with the highest relative abundance was selected to revise the annotated 5’ sequence (Figure S4A). The seed family assignments of the revised miRNAs were subsequently updated. Because the sequence revision could change miRNA suffixes and the ranking of *Hmi-novel-* miRNA, the names of all 545 miRNAs were reassigned accordingly (Figure S4A). Structural features, including mature length, mature base-pairing ratio, maximum bulge asymmetry, 3’ overhang, sequential apical loop size, as well as read counts, mature-strand per-base coverage, and annotations scores, were re-calculated (Figure S4B).

### Score Assignment

The score of each of the 545 predicted miRNA was determined by subtracting penalties and adding bonuses from an initial score of 100. The penalties were assigned based on structural features, including mature- and star- strand lengths, minimum free energy, mature-strand base-pairing ratio, maximum bulge asymmetry, 3’ overhang, structural and sequential apical loop sizes of the pre-miRNA (Figure S5A); expression features including total mature plus star read counts and maximum mature-strand per-base coverage (Figure S5B-C), and prediction quality metrics, including in-cluster ratio of mature and star strands, 5’ heterogeneity, and prediction algorithm (Figure S5D-F). Final scores exceeding 100 due to the bonus (prediction algorithm) were capped at 100.

The penalty thresholds for minimum free energy, mature base-pairing ratio, sequential and structural apical loop sizes were determined using all curated pre-miRNAs in all non-*H. miamia* bilaterian species in MirGeneDB 3.0 (12). Thresholds for maximum bulge asymmetry, which may cause a kink in the stem of the pre-miRNA, were based on previous structural analyses (29). Thresholds for mature- and star- strand lengths were based on MiRGeneDB recommendations (6, 98). Thresholds for 3’ overhang were determined from published biochemical studies (30–32). The penalty for mature + star abundance is determined according to the ranked percentile in cumulative abundance.

Annotation scores are provided in Column 6 of the GFF file, whereas the values of all individual scoring terms and the penalty are provided in Column 9 (Supplemental Data S2). Thresholds, penalty functions, and rationale for each scoring term are provided in Table S1. The criteria used to define the highest-confidence miRNAs are provided in Table S2.

Adjusted scores (Score.adj) used for the analyses in Figure 2L, were calculated by excluding penalties for expression features and prediction-quality metrics because equivalent raw data for the curated species were unavailable.

### *H. miamia* miRNA nomenclature

The 7mer seed sequences (g2-g8) of all the curated mature miRNAs in MirGeneDB v3.0 were extracted to generate a reference seed database. For each of the 545 annotated *H. miamia* miRNAs, the 7mer seed sequence of the dominant strand (mature strand) was extracted and searched against the reference database for exact matches. Reference miRNAs were queried sequentially in the following order of priority: (1) human; (2) conventional model organisms (*C. elegans, D. melanogaster, D. rerio, M. musculus*); (3) all other bilaterian species; (4) xenacoelomorph species; and (5) non-bilaterian metazoans. Matches identified from the dominant strand were assigned the value “7mer” for the *seed_match_type* flag in the annotation output. If no match was identified, the same search was performed using the star strand, and successful matches were assigned the value “7mer_alt” value. For these matched miRNAs, the 3’ non-seed region was subsequently compared only with curated miRNAs belonging to the first matched reference group using a customized R script based on pairwise alignment, and the curated miRNA with the highest alignment score was recorded in the *ref_mGDB* annotation field in the annotation output. The 7mer seed families for these miRNAs were referred to as curated families.

miRNA without a conserved 7mer seed were annotated as *H. miamia-*specific and assigned to HMI-NOVEL-n seed families. HMI-NOVEL-n families were ranked according to their cumulative read abundance and named sequentially based on the ranking. For miRNAs without strand bias as defined above (abundance of dominant strand less than twice the other strand), the seed sequences of both strands were evaluated. If the seed of the star strands matched a higher-ranked HMI-NOVEL family, the miRNA was assigned to that family.

Finally, alphabet suffixes were assigned for miRNAs sharing identical seed sequences (7mer seed for curated families, and 6mer seed for *H. miamia-*specific families) but differing in their non-seed sequences, whereas numeric suffixes were assigned to miRNAs with identical mature sequences. All naming information is provided in Table S4.

### miRNA expression dynamic analysis

To analyze miRNA expressions, miRNA reads were processed and filtered as described above. The filtered reads were then mapped to the *H. miamia* genome by star v2.7.10a using default parameters. Gene counting was done by featureCounts in subread v1.6.2 with reads mapped to multiple miRNA loci counted with *-M --fraction* options and reads per million (RPM) values of 5p, 3p, and the pre-miRNA loci were calculated using all reads mapped any miRNA loci as the normalizer. The fuzz c-mean clustering was performed by Mfuzz v2.62.0 with *c = 4, m = 1.35* options, using the sum RPM of 5p and 3p strands (45). The early cleavage stage was excluded from this step to increase the clustering quality but included in subsequent analyses. The differential expression analysis of sRNA seq from anterior and posterior fragment samples was performed by DESeq2 with default parameters (99). Significantly differentially expressed genes indicate those with |FC| >1.5 and p.adj < 0.05 in DESeq2.

### Bulk ATAC seq data analysis

Filtered and demultiplexed paired-end bulk ATAC-seq reads from *H. miamia* embryonic developmental stages (NCBI PRJNA1167694) and the 0-hour post-amputation anterior and posterior wound samples (NCBI PRJNA512373) were mapped to the PacBio using bowtie2 v2.5.0 and the alignment was sorted by Samtools v1.16.1 (92). CPM-normalized coverage track for genome browser visualization is generated using the bamcoverage package of deeptools v3.5.6 (100).

### RT-PCR for *hmi-let-7/hmi-mir-125-a* primary transcript

Total RNA from the PL, EJ and SMA developmental stages of *H. miamia* and mixed stage *C. elegans* was treated with Turbo DNase (ThermoFisher Cat: AM2239) to remove genomic DNA and purified using Zymo RNA Clean & Concentrate 5 kit (Cat: R1013). RNA concentration was determined using Qubit RNA BR assay kit (ThermoFisher, Cat# Q10210). First strand cDNA was synthesized from 450 ng of total RNA using CoWin HifiScript RT mastermix (Cat: CW3371) with a mixture of random hexamer and oligo dT. Genomic DNA from mixed stage *H. miamia* was extracted using NEB Monarch Spin gDNA Extraction Kit (Cat: T3010S). The PCR amplification was performed using CoWin Hi-sincere PCR mix (Cat: CW3419) using primers GAGGTAGTGTGATAAACAGTTTTGAAC and CATGGGCTTAGGTCTCAGGGAC for 35 cycles.

### Evolutionary analysis of intron in LET-7/MIR-125/MIR-100 clusters

Genomic coordinates of all LET-7/MIR-125/MIR-100 family miRNAs were obtained from MirGeneDB v3.0 except for those from *H. miamia*. Genomic clusters were defined as two or more family members located on the same strand within a genomic span of less than 10 kb. For each cluster, the genomic interval between the two outermost miRNAs was extracted from the corresponding reference genome from Ensembl Metazoa release 59 or NCBI genomes. Splice donor and acceptor sites were predicted using ASSP MultiSEQ using either default score thresholds (false splice acceptor sites ≥2.2 and false splice donor sites ≥4.5; Figure 5E) or the most stringent thresholds available for the software (false splice site ≥7.0 and false splice donor sites ≥9.0; Figure S12B) (70). Only constitutive splice donor and accepter sites were retained and analyzed.

### *H. miamia* miRNA Target prediction

Target prediction was performed as described before (37, 57). The sites with perfect seed pairing were defined as perfect Watson-Crick base pairing at g2-g7. The 3’ compensatory sites with imperfect seed and 3’ pairing were defined as a seed pairing with one GU pair or a single nucleotide target-strand bulge at g5-g8, combined with at least three consecutive Watson-Crick base pairings at g11-g16 .

### *C. elegans* miRNA target analysis

The *C. elegans* miRNAs and target sites were used as a reference to test the quality of *H. miamia* target site predictions in this study. The *C. elegans* miRNA sequences and 3’ UTR sequences were obtained from *WBCel279* (101, 102). The *C. elegans* miRNA target sites predictions were obtained from TargetScan 6.2 (103).

## ACKNOWLEDGEMENT

We thank Harvard Bauer Core for the PacBio sequencing library preparation and sequencing. We thank Adam Freedman from the Harvard FAS Bioinformatics Group for the help of genome annotation. We thank members of the Srivastava Lab, Ambros Lab, and Veksler-Lublinsky Lab for the discussions that helped advance this project. We thank Amber Rock and Paul Bump for helping with embryo manipulation, Kaitlyn Loubet-Senear for helping with bioinformatics, and D. Marcela Bolaños for helping with animal culturing and amputation. We thank Bastian Fromm for commenting on the annotation evaluation. We thank Alejandro Vasquez-Rifo, Luisa Cochella, Catriona Breen , Kaitlyn Loubet-Senear and Carlos Rivera-López for commenting on this manuscript.

## DATA AVAILABILITY

The PacBio genome and miRNA sequencing raw data will be available at Gene Expression Omnibus and NCBI genome. Key source codes will be deposit at GitHub

## AUTHOR CONTRIBUTIONS

Y.D., V.A., M.S. designed research; Y.D., T.S., I.V.L. performed research; Y.D., T.S., D.E.K., I.V.L. analyzed data; and Y.D., V.A., and M.S. wrote the paper.

