## Supplemental Files for "A comprehensive resource for studying microRNA evolution and microRNA-mediated development and whole-body regeneration in the acoel worm *Hofstenia miamia*"

#### SUPPLEMENTAL DATASETS AND TABLES

**Dataset S1. GTF files for *H. miamia* annotations, including transcripts and miRNAs.**

**Dataset S2. GTF file for the *H. miamia* miRNA annotation in this study.** The annotation file contains coordination of all 545 *H. miamia* pre-miRNA, 5p, and 3p features. The scores are provided in Column 6. All structural, expression values used to generate the annotation scores are provided in Column 9.

**Table S1. Thresholds and rationales for the annotation score penalty/bonus terms.**

**Table S2. Thresholds and rationales to determine the highest-confidence miRNAs.**

**Table S3. Summary of sequence, structure, expression, and prediction quality features of predicted *H. miamia* miRNA loci. A.** All the 545 *H. miamia* miRNA loci. **B.** 154 highest-confidence miRNA loci.

**Table S4. Summary of the nomenclature and evolutionary node of origin of all nominated 545 *H. miamia* miRNAs**

**Table S5. Results of *H. miamia* sRNA-seq for development profiling. A-B.** Raw and normalized read counts (Reads per Million, RPM) of the 5p and 3p strands from all 545 *H. miamia* miRNA loci across 11 developmental stages, calculated as the mean of three biological replicates per developmental stage. **C-D.** Raw and normalized read counts (Reads per Million, RPM) of the 5p and 3p strands from all 545 *H. miamia* miRNA loci across 11 developmental stages, with each biological replicate shown individually. **E-J.** Raw and normalized read counts of the mature strands, star strands, and mature+star strands. Non-integer values resulting from the *-M --fraction* option in *featureCounts* were rounded to the nearest integer.

**Table S6. Results of target prediction for the highest-confidence miRNAs. A.** Predicted miRNA::target interaction pairs and information of the binding sites with perfect seed pairing configuration. **B.** Predicted miRNA::target interaction pairs with 3' complementary pairing configuration. In this category, only GU wobble at g5-g8 or target strand single-nucleotide bulge at g5-g7 in the seed region pairing, accompanied with at least 3 consecutive Watson-Crick pairings at g11-g16 region, were included according to previous genetic study (1).

**Table S7. Differential expression analysis results of the sRNA seq for anterior and posterior fragments. A.** Differential expression analysis results comparing the miRNA expression in the anterior fragment (H00) and posterior fragment (T00). Fold change (FC) represents the ratio of expression in the anterior fragment to the posterior fragment. **B-C.** RPM and raw reads for the three replicates in anterior and posterior fragments, values were aggregated per pre-miRNA strands. **D.** Raw reads for the three replicates in anterior and posterior fragments, split by miRNA 5p and 3p. Non-integer values in C and D were rounded as shown in Table S5D.

### Table S1 Penalty thresholds for the Annotation Score

| | Term | Threshold | Penalty | Rationale |
|---|---|---|---|---|
| 1 | **Mature Length (Lm)** | 20 ~ 26 | 0 | Cutoff by miRGeneDB (mGDB) |
| | | = 19 | 5 | Outside mGDB cutoff; above 99 rank percentile of miRBase curation |
| | | = 18 | 10 | Outside mGDB cutoff; above 99.9 rank percentile of mB curation |
| 1 | **Star Length (Ls)** | 20 ~ 26 | 0 | mGDB curation |
| | | ≤19 | 5 | Outside mGDB range |
| 1 | **Minimum Free Energy (MFE)** | ≤ -17.4 kcal | 0 | Above 95 rank percentile of all mGDB curation |
| | | -17.4 ~ -14.7 kcal | 1-39 | Above 95 rank percentile of all mGDB curation |
| | | ≥ -14.7 kcal | 40 | Below 99 rank percentile of all mGDB curation |
| 1 | **Mature Base-Paring Ratio** | ≥ 0.69 | 0 | Above 95 rank percentile of all mGDB curation |
| | | 0.62 ~ 0.69 | 5-30 | Between 95-99 rank percentile of all mGDB curation |
| | | < 0.62 | 40 | Below 99 rank percentile of all mGDB curation |
| 1 | **Max Bulge Asymmetry** | ≤ 2 nt | 0 | No helix distortion |
| | | 3 nt | 15 | Helix distortion |
| | | 4 nt | 30 | Major Helix distortion; 99 rank percentile of all mGDB |
| | | ≥ 5 nt | 40 | Major Helix distortion; below 99 rank percentile of all mGDB |
| 1 | **3' Overhang** | 2 nt | 0 | No Dicer processing compromise |
| | | 3 nt | 5 | Minor Dicer processing compromise (Lee et al. 2022) |
| | | 1 nt | 10 | Moderate Dicer processing compromise (Lee et al. 2022)) |
| | | 0, -1, or ≥ 4 nt | 15 | Putative major Dicer processing compromise (lacking biochemical validation) |
| | | ≤ -2 nt | 40 | Incompatible with Dicer processing (Feng et al. 2012) |
| 1 | **Structural Apical Loop Size** | ≥ 6 nt | 0 | Optimal length for Drosha and Dicer processing (Feng et al. 2012) |
| | | 5 nt | 3 | Minor processing compromise (Feng et al. 2012) |
| | | 4 nt | 5 | Moderate processing compromise (Feng et al. 2012) |
| | | 3 nt | 10 | Min. RNA stem-loop; major processing compromise (Feng et al. 2012) |
| 1 | **Sequential Loop Size** | ≥ 6 nt | 0 | Above 95 rank percentile of all mGDB curation |
| | | 5 nt | 5 | Between 95-99 rank percentile of all mGDB curation |
| | | <= 4 nt | 10 | Below 99 rank percentile of all mGDB curation |
| 2 | **Total Mature+Star Reads (total m+s)** | > 1995 | 0 | Ranked cumulative counts account for 99.9% of all reads |
| | | 270 ~ 1995 | 1~24 | Ranked cumulative counts account for 0.1%-0.01% of all reads |
| | | < 270 | 25 | Ranked cumulative counts account for < 0.01% of all reads |
| 2 | **Max Mature per-Base coverage (m_pBC)** | > 6.1X | 0 | Rank percentile above 50% |
| | | 6.1 ~ 0.22X | 1-24 | Rank percentile between 90% and 50% |
| | | < 0.22X | 25 | Rank percentile below 90% |
| 3 | **5' heterogeneity of mature strand** | NA | 10 | |
| | | < 10% | 0 | |
| | | 10% ~ 50% | 1-39 | |
| | | >50% | 40 | |
| 3 | **In-cluster ratio of mature strand** | > 99% | 0 | |
| | | 99% ~ 75% | 1-39 | |
| | | < 75% | 40 | |
| 3 | **Detecting algorithm** | By both | -20 | |
| | | By one algorithm | 0 | |

**Filter Categories:**

1. Precursor/miRNA structural properties
2. Expression abundance
3. Prediction quality

### Table S2. Thresholds for highest-confidence miRNAs

| 0 | Term | Threshold | Rationale |
|---|---|---|---|
| 1 | **Mature Length (Lm)** | ≥ 20 | Cutoff by miRGeneDB (mGDB) |
| 1 | **Star Length (Ls)** | ≥ 20 | Cutoff by miRGeneDB (mGDB) |
| 1 | **Minimum Free Energy (MFE)** | ≤ -14.7 kcal | Above 99 rank percentile of all mGDB curation |
| 1 | **Mature Base-Paring Ratio** | ≥ 0.62 | Above 99 rank percentile of all mGDB curation |
| 1 | **Max Bulge Asymmetry** | ≤ 3 nt | Without major helix distortion |
| 1 | **3' Overhang** | 0 ~3 nt | No major Dicer processing compromise |
| 1 | **Structural Apical Loop Size** | ≥ 3 nt | Minimum RNA stem-loop; major processing compromise (Feng et al. 2012) |
| 1 | **Sequential Loop Size** | ≥ 6 nt | Above 99 rank percentile of all mGDB curation |
| 2 | **Total Mature + Star Reads** | > 1995 | Ranked cumulative counts account for 99.9% of all reads |
| 2 | **Max Mature per-Base coverage )** | > 6.1X | Rank percentile above 50% |
| 3 | **5' heterogeneity of mature strand** | < 20% | Arbitrary threshold |
| 3 | **In-cluster ratio of 3p + 5p strands** | ≥ 90% | Arbitrary threshold |

**Filter Categories:**

1. Precursor/miRNA structural properties
2. Expression abundance
3. Prediction quality

Figure S1

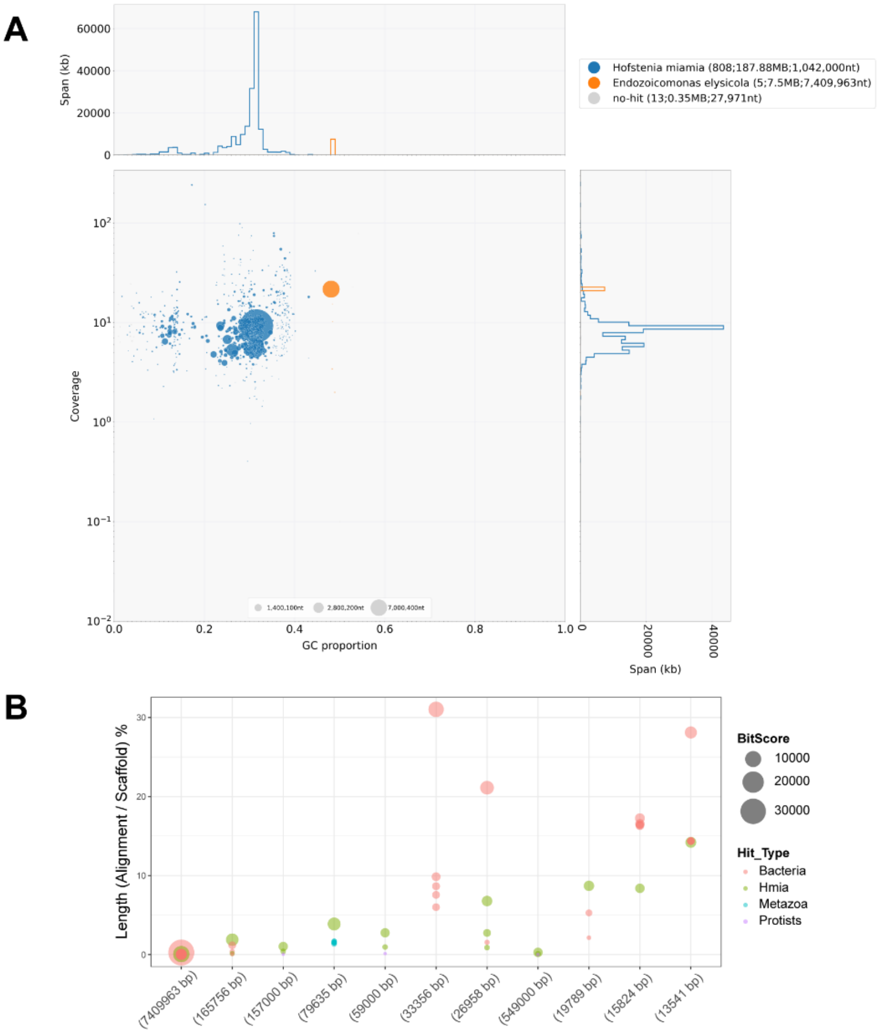


**C**

| | Eukaryota_OBD12 | Metazoa_ODB12 |
|---|---|---|
| Complete BUSCOs | 120 (93%) | 587 (87.4%) |
| Complete and single-copy BUSCOs | 61 (47.3%) | 296 (44.0%) |
| Complete and duplicated BUSCOs | 59 (45.7%) | 291 (43.3%) |
| Fragmented BUSCOs | 5 (3.9%) | 41 (6.1%) |
| Missing BUSCOs | 4 (3.1%) | 44 (6.5%) |
| Total BUSCO groups searched | 129 | 672 |

**Figure S1. Summary of *H. miamia* PacBio genome decontamination. A.** Blob plot of the *H. miamia* PacBio genome assembly before decontamination. The x-axis shows the GC content of each scaffold, and the y-axis shows the sequencing coverage. The color of each point indicates the taxonomic assignment of the top BLAST hit, and the size of each point is proportional to the scaffold length. **B.** The proportions of aligned sequences relative to the full scaffold length for the scaffolds whose top five BLAST hits were not *H. miamia.* To identify the potential contaminant scaffolds, all initially assembled PacBio scaffolds were searched against all available metazoan, protist, plants, and bacterial genomes using MegaBLAST. Scaffolds for which the top five alignment are not to the previously published *H. miamia* genome were classified as putative contaminants. Point size indicates the BLAST bit score, and the point color indicates the taxonomic group of the hit, with non-*H. miamia* hits grouped by kingdom. **C.** BUSCO assessment results of the current *H. miamia* annotation using the Eukaryota_OBD12 and Metazoa_ODB12 reference datasets. The exported transcriptome for BUSCO assessment includes all isoforms for each gene, resulting in high proportion of duplicated BUSCOs.

Figure S2

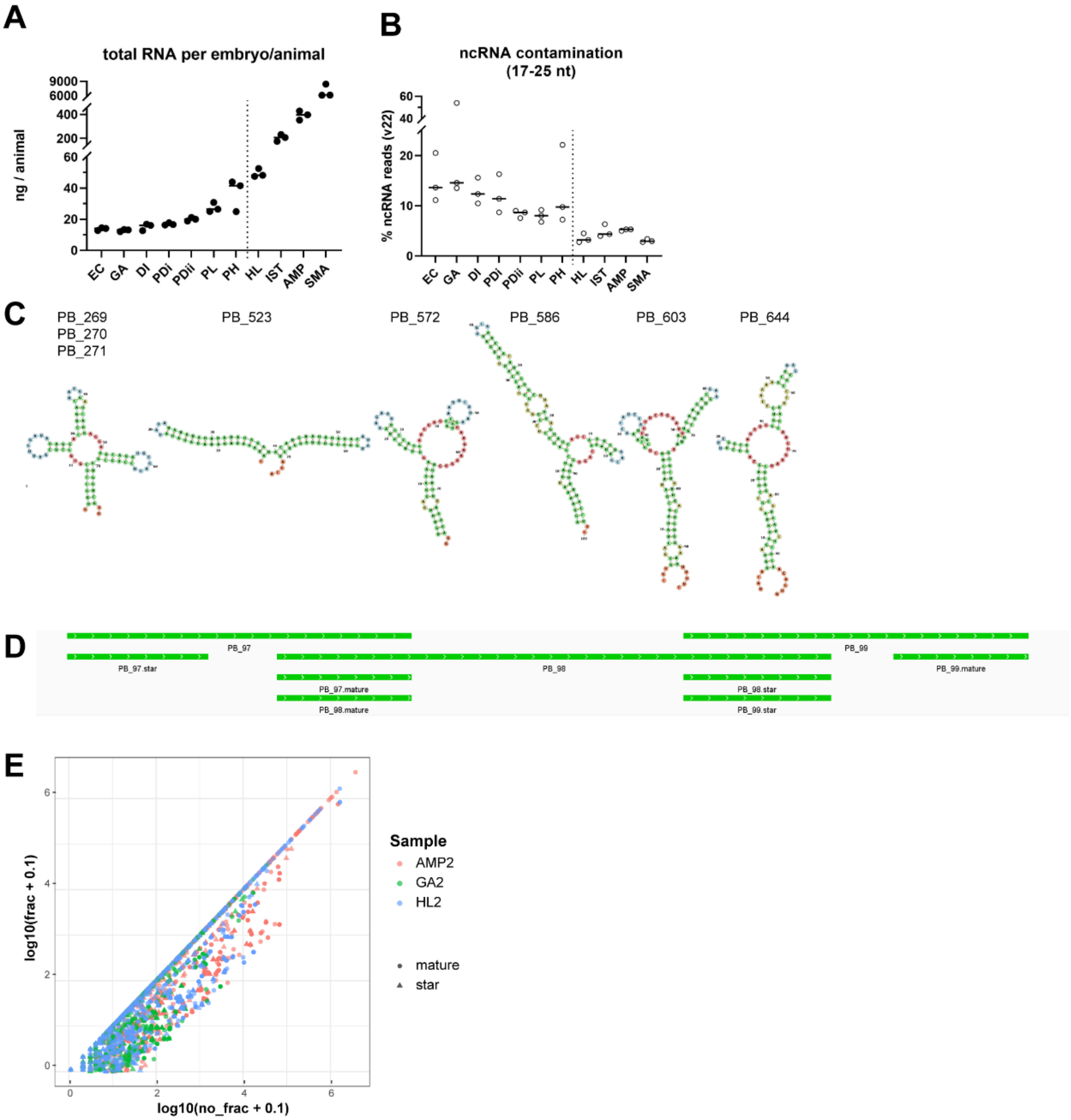


**Figure S2. sRNA sequencing and false positive predictions for the *H. miamia* miRNA annotation.** **A.** The total RNA yield per embryo or animal for each developmental stage. The dashed line separates embryonic and post-embryonic development. **B.** Proportion of contaminating non-coding RNAs (17-25 nt) among sRNA-seq reads at each developmental time point. The non-coding RNA sequences for *H. miamia* were obtained from RNAcentral v22. **C.** Secondary structures of the predicted pre-miRNAs with nonclassical hairpin structures. PB_269/270/271 have identical sequences. Secondary structures were predicted by FORNA (2). **D.** Genomic coordinates of PB_97/98/99. Note that PB_98 was an artificial prediction generated from the mature strand of PB_97 and the star strand of PB_99 and was therefore filtered out. **E.** Scatter plot comparing miRNA read counts using fractional (y-axis) and non-fractional (x-axis) counting approaches. Read counts were shown as log10-transformed after adding pseudocount of 0.1 to each locus. Three representative libraries are shown: AMP2 (highest total reads), GA2 (lowest total reads), and HL2 (highest mapping rate).

Figure S3

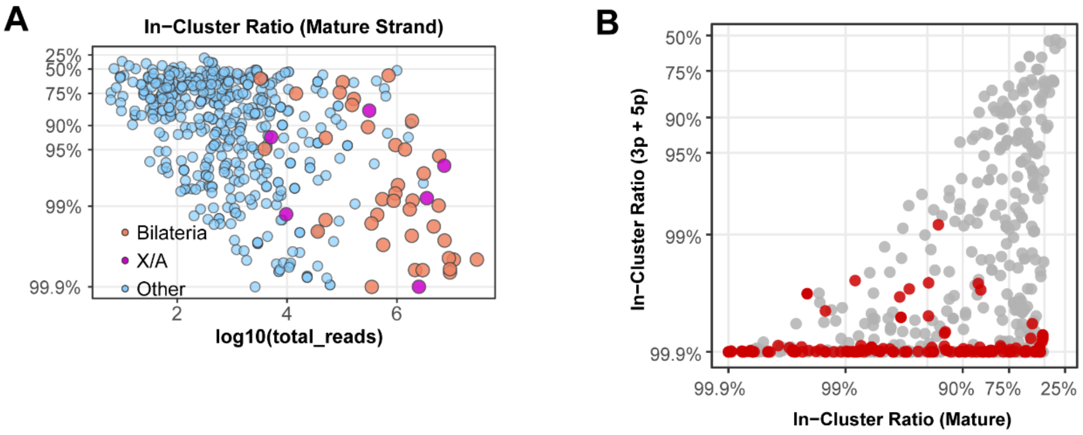


**Figure S3. Comparison of the in-cluster ratio calculation methods. A.** Scatter plot of the $\log_{10}$ in-cluster ratio (y-axis) against the $\log_{10}$ total read (x-axis) of all nominated *H. miamia* miRNAs shown for comparison with **Figure 2H**. miRNAs are colored according to the evolutionary node of origin of their seed families. The in-cluster ratio was calculated as the proportion of reads mapping to the mature strand relative to all reads mapping to the pre-miRNA locus with ±10 bp flanking regions (3). **B.** Scatter plot comparing the in-cluster ratios that were calculated using combined 3p and 5p strands (y-axis; Figure 2G) and using mature strand only (x-axis; Figure S3A), red points indicate highest-confidence miRNAs.

#### Figure S4

**A**

log2FC iso/ref (scale: 3, 6, 9, 12) — log10_isomiR_count (scale: 2, 3, 4, 5)

| isomiR | log10_isomiR_count | log2FC iso/ref | global_isomiR_% | top_stage_log2FC | #_stages_with_higher_isomiR | longest_dev_stages_streak | seed_with_higher_node | detected_hi_conf_seed | REVISED | Ref_family (node of origin) | | Revised_family (node of origin) | |
|---|---|---|---|---|---|---|---|---|---|---|---|---|---|
| PB_332-iso-p3 | | 3.16 | 37.6 | 15.95 | 5 | 5 | | | ◆ | mir-2704 | C. teleta | hmi-novel-2 | H.miamia |
| PB_332-iso-p2 | | 2.84 | 30.2 | 15.82 | 4 | 3 | | | | mir-2704 | C. teleta | | |
| PB_509-iso-p3 | | 0.95 | 41.4 | 1.62 | 1 | 1 | | | ◆ | hmi-novel-3 | H.miamia | hmi-novel-4 | H.miamia |
| PB_332-iso-p1 | | 2.02 | 17 | 14.91 | 3 | 3 | ■ | | | mir-2704 | C. teleta | | |
| PB_302-iso-m2 | | 1.34 | 71.6 | 0.34 | 5 | 4 | ■ | ■ | ◆ | hmi-novel-8 | H.miamia | mir-335 | Eutheria |
| PB_332-iso-m1 | | 1.15 | 9.3 | 13.6 | 4 | 4 | | | | mir-2704 | C. teleta | | |
| PB_455-iso-p2 | | 6.06 | 82.6 | 6.33 | 8 | 6 | | | ◆ | hmi-novel-14 | H.miamia | hmi-novel-32 | H.miamia |
| PB_241-iso-p1 | | 7.1 | 89.1 | 7 | 7 | 4 | | ■ | ◆ | mir-4060 | C. intestinalis | mir-2056 | Branchiostoma |
| PB_74-iso-p2 | | 7.32 | 97.8 | 7.78 | 4 | 2 | | | ◆ | hmi-novel-103 | H.miamia | hmi-novel-6 | H.miamia |
| PB_352-iso-m3 | | 7.54 | 99.2 | 7.06 | 6 | 4 | | ■ | ◆ | hmi-novel-43 | H.miamia | hmi-novel-44 | H.miamia |
| PB_252-iso-m3 | | 12.26 | 100 | 9.69 | 11 | 11 | | | ◆ | hmi-novel-59 | H.miamia | hmi-novel-61 | H.miamia |
| PB_455-iso-p3 | | 3.49 | 13.8 | 3.68 | 3 | 1 | | | | hmi-novel-14 | H.miamia | | |
| PB_484-iso-p2 | | 1.64 | 73.5 | 2.02 | 1 | 1 | | ■ | ◆ | hmi-novel-74 | H.miamia | hmi-novel-9 | H.miamia |
| PB_493-iso-m1 | | 3.54 | 65 | 2.28 | 2 | 1 | | | ◆ | hmi-novel-42 | H.miamia | hmi-novel-49 | H.miamia |
| PB_241-iso-m1 | | 3.96 | 10.1 | 3.92 | 3 | 3 | | | | mir-4060 | C. intestinalis | | |
| PB_263-iso-m2 | | 0.62 | 59.6 | 0.39 | 1 | 1 | ■ | ■ | ◆ | hmi-novel-8 | H.miamia | mir-205 | Vertebrata |
| PB_111-iso-m2 | | 2.2 | 82 | 3.01 | 1 | 1 | | ■ | ◆ | hmi-novel-8 | H.miamia | hmi-novel-9 | H.miamia |
| PB_512-iso-m2 | | 2.87 | 86.5 | 2.39 | 1 | 1 | ■ | ■ | ◆ | hmi-novel-8 | H.miamia | mir-205 | Vertebrata |
| PB_493-iso-p1 | | 2.12 | 24.3 | 4.63 | 2 | 1 | | | | hmi-novel-42 | H.miamia | | |
| PB_260-iso-p2 | | 2.54 | 80.9 | 2.39 | 1 | 1 | | | ◆ | hmi-novel-5 | H.miamia | hmi-novel-6 | H.miamia |
| PB_326-iso-m2 | | 2.33 | 82.9 | 3.22 | 1 | 1 | | | ◆ | hmi-novel-102 | H.miamia | hmi-novel-102 | H.miamia |
| PB_490-iso-p2 | | 2.59 | 79.9 | 2.71 | 2 | 1 | | ■ | ◆ | mir-335 | Eutheria | hmi-novel-9 | H.miamia |
| PB_437-iso-m2 | | 2.19 | 69.7 | 3.15 | 1 | 1 | | | ◆ | hmi-novel-80 | H.miamia | mir-1401 | O. anatinus |
| PB_545-iso-p2 | | 2.36 | 77 | 2.85 | 0 | 0 | | ■ | ◆ | mir-335 | Eutheria | hmi-novel-9 | H.miamia |
| PB_403-iso-m2 | | 1.98 | 78.9 | 2.55 | 1 | 1 | ■ | ■ | ◆ | hmi-novel-8 | H.miamia | hmi-novel-9 | H.miamia |
| PB_549-iso-m2 | | 2.11 | 74.5 | 3.17 | 0 | 0 | | | ◆ | hmi-novel-123 | H.miamia | hmi-novel-122 | H.miamia |
| PB_78-iso-m2 | | 7.78 | 90.5 | 5.91 | 5 | 3 | | | ◆ | hmi-novel-100 | H.miamia | hmi-novel-100 | H.miamia |
| PB_2-iso-m2 | | 2.62 | 85.6 | 2.92 | 0 | 0 | ■ | ■ | ◆ | hvu-novel-12 | H. vulgaris | mir-205 | Vertebrata |
| PB_30-iso-p3 | | 5.31 | 83.7 | 5.55 | 0 | 0 | | | ◆ | hmi-novel-77 | H.miamia | hmi-novel-78 | H.miamia |
| PB_409-iso-m2 | | 4.83 | 97.4 | 4.25 | 1 | 1 | ■ | ■ | ◆ | hmi-novel-8 | H.miamia | hmi-novel-9 | H.miamia |
| PB_67-iso-p2 | | 3.81 | 36.9 | 4.7 | 0 | 0 | | | ◆ | hmi-novel-125 | H.miamia | hmi-novel-124 | H.miamia |

**B**

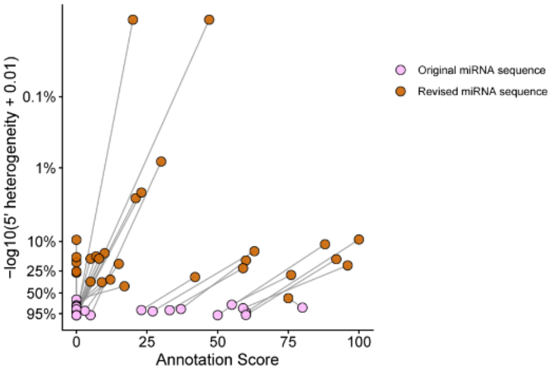


**Figure S4. Revision of miRNA sequences for 5' heterogeneous miRNAs. A.** Summary of the 5' heterogeneity and sequence revision outcomes of 5' heterogeneous miRNAs. **The left panel** lists the isomiR ID as described in the Methods. **The first (gray) column** of the heatmap shows the $\log_{10}$ total counts of the isomiR, calculated using the counting algorithm described in the Methods. **The second (grey) column** shows the $\log_2$ fold change in read counts between the isomiR and the reference miRNA (the miRNA sequence predicted by miRDeep2 and/or sRNAbench). **The third (orange) column** shows the global proportion of reads corresponding to the isomiR within the 5' heterogeneous miRNA locus (defined as the miRDeep2/sRNAbench output 5' coordinate with ±3 bp flanking), calculated by combining all libraries. Color labeling indicates that the isomiR accounts for more than 50% of reads in the 5' heterogeneous locus globally. **The fourth (green) column** shows the highest $\log_2$ fold change between the isomiR count with the reference miRNA count across all developmental stages. Color labeling indicates that at this stage the isomiR count is at least two-fold higher than the reference miRNA counts in at least two of the three biological replicates. **The fifth (blue) column** shows the number of developmental stages in which the isomiR count exceeds the reference miRNA count in at least two of the three biological replicates. Color labeling indicates that the isomiR exceeds the reference miRNA in more than half (6) of the developmental stages. **The sixth (blue) column** shows the longest consecutive streak of developmental stages in which the isomiR count exceeds the reference miRNA count in at least two of the three biological replicates. Color labeling indicates the longest streak is greater than three developmental stages. **The seventh (purple) column** indicates whether sequence revision reassigns the miRNA to a more evolutionarily conserved seed family. Color labeling indicates positive cases. **The eight (blue) column** indicates whether sequence revision reassigns the miRNA to a seed family that was included in the highest-confidence miRNA list. Color labeling indicates positive cases. Brown circled diamonds indicate that miRNAs whose 5' sequences were revised to the corresponding isomiR sequences. Node of family origin assignments were obtained from MirGeneDB v3.0. **B.** Comparison of the $\log_{10}$ 5' heterogeneity (y-axis) and annotation score (x-axis) between the original and revised miRNA sequences. The revised annotation scores were calculated using updated feature quantification and penalties for mature length, maximum bulge asymmetry, mature base-pairing ratio, 3' overhang, sequential apical loop size, reads counts, and maximum per-base coverage. For each miRNA locus, the points of original (pink) and revised sequences (orange) are connected by a line.

Figure S5

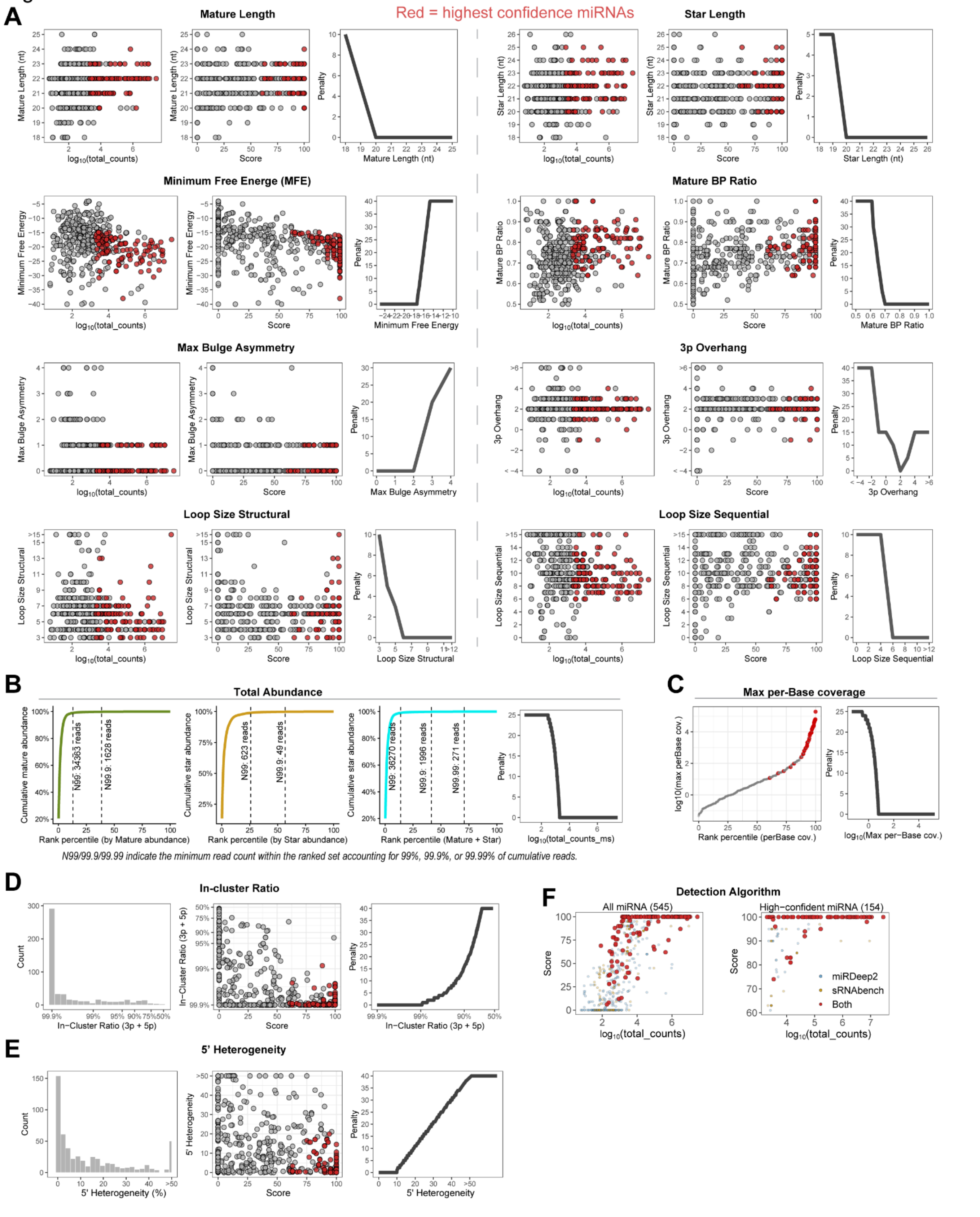


**Figure S5. Components of the annotation score of *H. miamia* miRNAs. A.** Quantification of structural features, including mature length, star length, minimum free energy, mature base-pairing ratio, maximum bulge asymmetry, 3’ overhang, structural apical loop size, and sequential apical loop size. For each feature, the left panels show the relationship of quantification of the feature (y-axis) and $\log_{10}$ total read counts (x-axis), the middle panels show the relationship of quantification of the feature (y-axis) and annotation score (x-axis), the right panels show the relationship of quantification of the features (x-axis) and assigned penalty (y-axis). Red points indicate highest-confidence miRNA loci. **B.** Summary of the total abundance. The left three panels show the cumulative abundance against rank percentile of the mature (green), star (brown) and mature + star (cyan) reads. Dashed vertical lines indicate the rank percentile corresponding to cumulative abundance account for 99%, 99.9% and 99.99% of all mapped reads, with the corresponding minimum reads labeled. The right panel shows the relationship between $\log_{10}$ of cumulative mature + star abundance (x-axis) and the assigned penalty. **C.** Summary of the maximum per-base coverage of all predicted miRNAs among all libraries. The left panel shows $\log_{10}$ maximum per-base coverage against rank percentile. The right panel shows the relationship between $\log_{10}$ maximum per-base coverage (x-axis) and the assigned penalty (y-axis). Per-base coverage of a miRNA locus was calculated as the total number of sRNA-seq reads mapped to the mature strand divided by the revised mature strand length after the 5’ heterogeneity revision. **D-E.** Quantification of the in-cluster ratio calculated using combined 3p and 5p reads (**D**) and 5’ heterogeneity (**E**). For each feature, the left panels show the distribution of the quantified values, the middle panels show the relationship between the quantified feature (y-axis) and the annotation score (x-axis), and the right panels show the relationship between the quantified features (x-axis) and the assigned penalty (y-axis). **F.** Distribution of the detection algorithm of all *H. miamia* miRNAs (left) and the highest-confidence miRNAs (right) on a scatter plot of $\log_{10}$ total counts (x-axis) and annotation score (y-axis). Colors indicate the prediction algorithm(s) that identified each miRNA locus (miRDeep2, sRNAbench, or both).

Figure S6

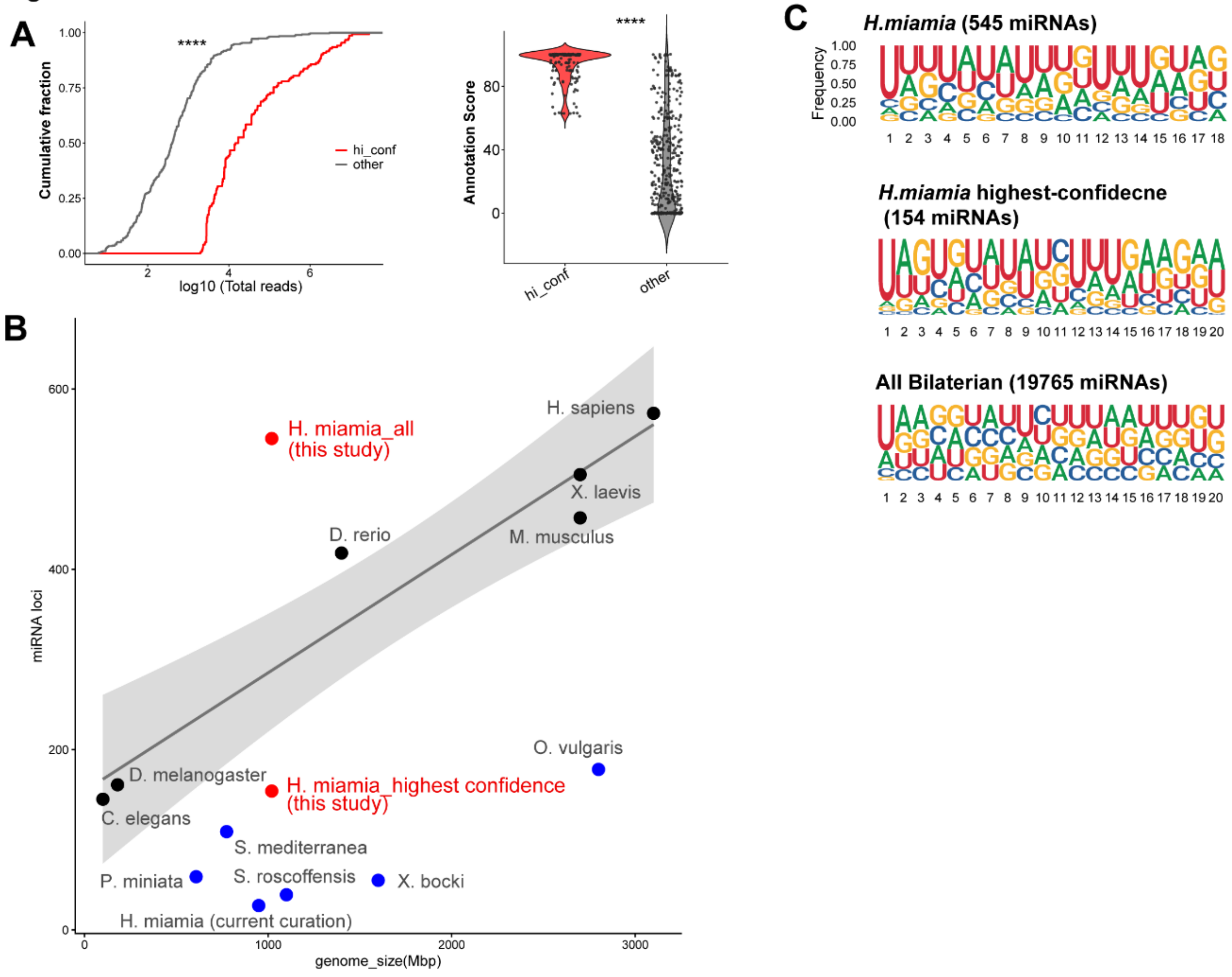


**Figure S6. Comparison between the highest-confidence miRNA subset and the remaining annotated miRNAs. A.** (Left) Empirical cumulative distribution of log10-transformed total reads comparing highest-confidence miRNAs and the remaining annotated miRNAs. Statistical significance was assessed using a two-sample K-S test. (Right) Violin plot comparing annotation scores between highest-confidence miRNAs and the remaining annotated miRNAs. Statistical significance was assessed using a two-sided Wilcoxon rank-sum test. ****, $P < 0.0001$. **B.** Relationship between annotated miRNA loci and genome size across species. The number of annotated miRNA loci and genome size for each species were obtained from MirGeneDB 3.0. The solid grey line represents the linear regression fit, and the shaded area indicates the 95% confidence interval. **C.** Nucleotide composition of all *H. miamia* miRNAs (top), highest-confidence miRNAs (center) and all curated bilaterian miRNAs (bottom). For *H. miamia,* only the dominant strand (5p or 3p) with higher total read counts was analyzed. Sequence logos were generated by ggseqlogo (4).

Figure S7

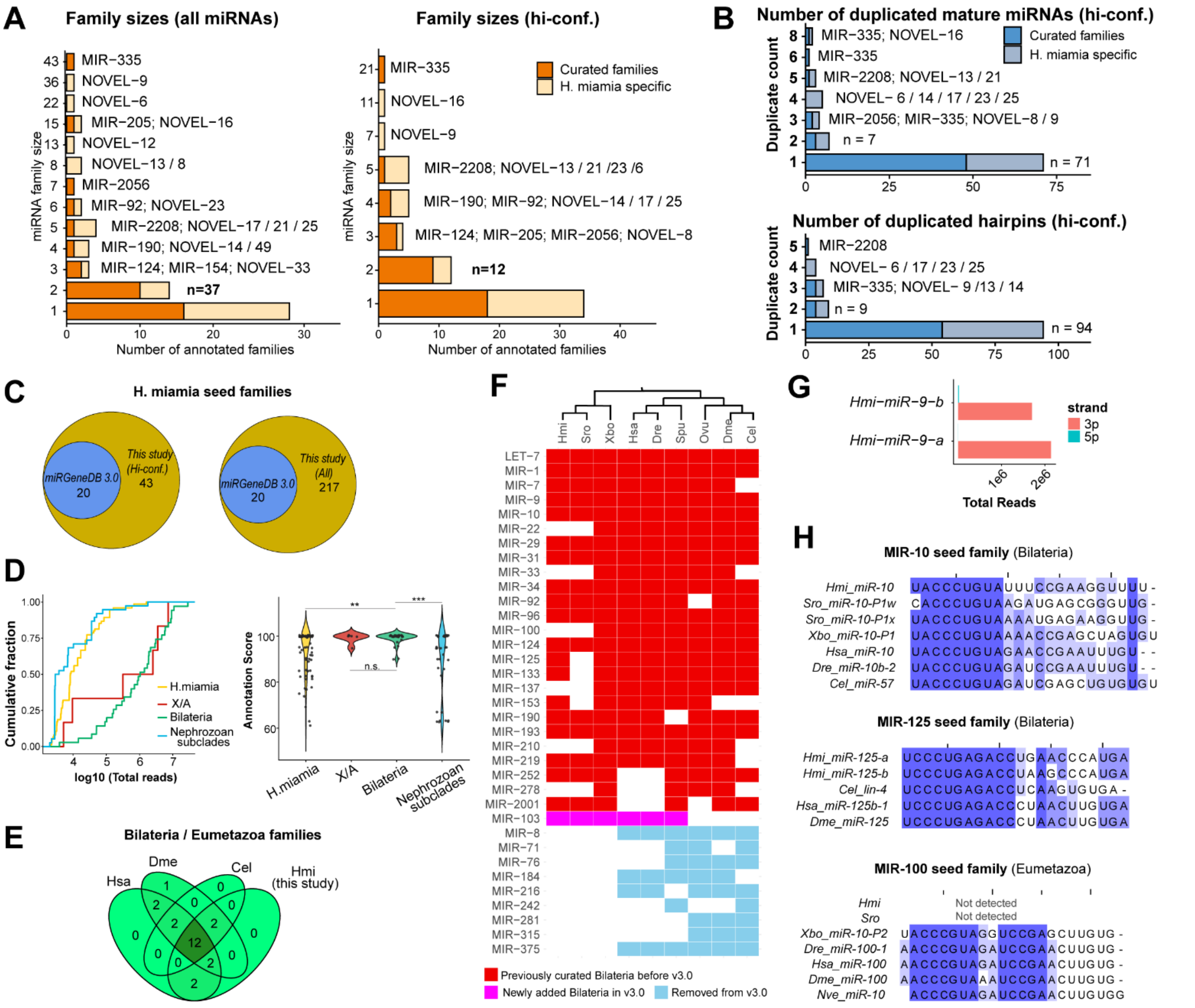


**Figure S7. Supplemental results for the evolutionary analysis of *H. miamia* miRNAs. A.** Distributions of the *H. miamia* seed family sizes. Curated families denote families with evolutionary node-of-origin curation that is not *H. miamia* . **B.** Distribution of the numbers of identical mature miRNA sequences (top) and identical pre-miRNA sequences (bottom). **C**. Venn diagrams showing *H. miamia* seed families previously curated in MirGeneDB v3.0 and that identified in this study for all miRNAs (right) and highest-confidence miRNAs (left). **D.** (Left) Empirical cumulative distribution of log10-transformed total read counts of all miRNAs categorized by the evolutionary node of origin of their seed families. X/A, Xenacoelomorpha/ Acoela. (Right) Violin plots comparing annotation scores among all miRNAs grouped by the evolutionary node of origin of their seed families. Significance was assessed using a two-sided Wilcoxon rank-sum test with BH correction. **, $p < 0.01$; **** $p < 0.0001$, n.s., not significant with $p > 0.05$. **E.** Venn diagram showing the overlap of bilaterian/eumetazoan seed families between *H. miamia* in this study and in human (Hsa), fruit fly (Dme), C. elegans (Cel). **F.** Presence or absence of all bilaterian seed families in *H. miamia* and representative bilaterian species. **G.** Total read counts of the 3p and 5p strands of MIR-9 family miRNAs. **H.** Sequence alignments of the mature strands of MIR-10, MIR-125, and MIR-100 seed families.

Figure S8

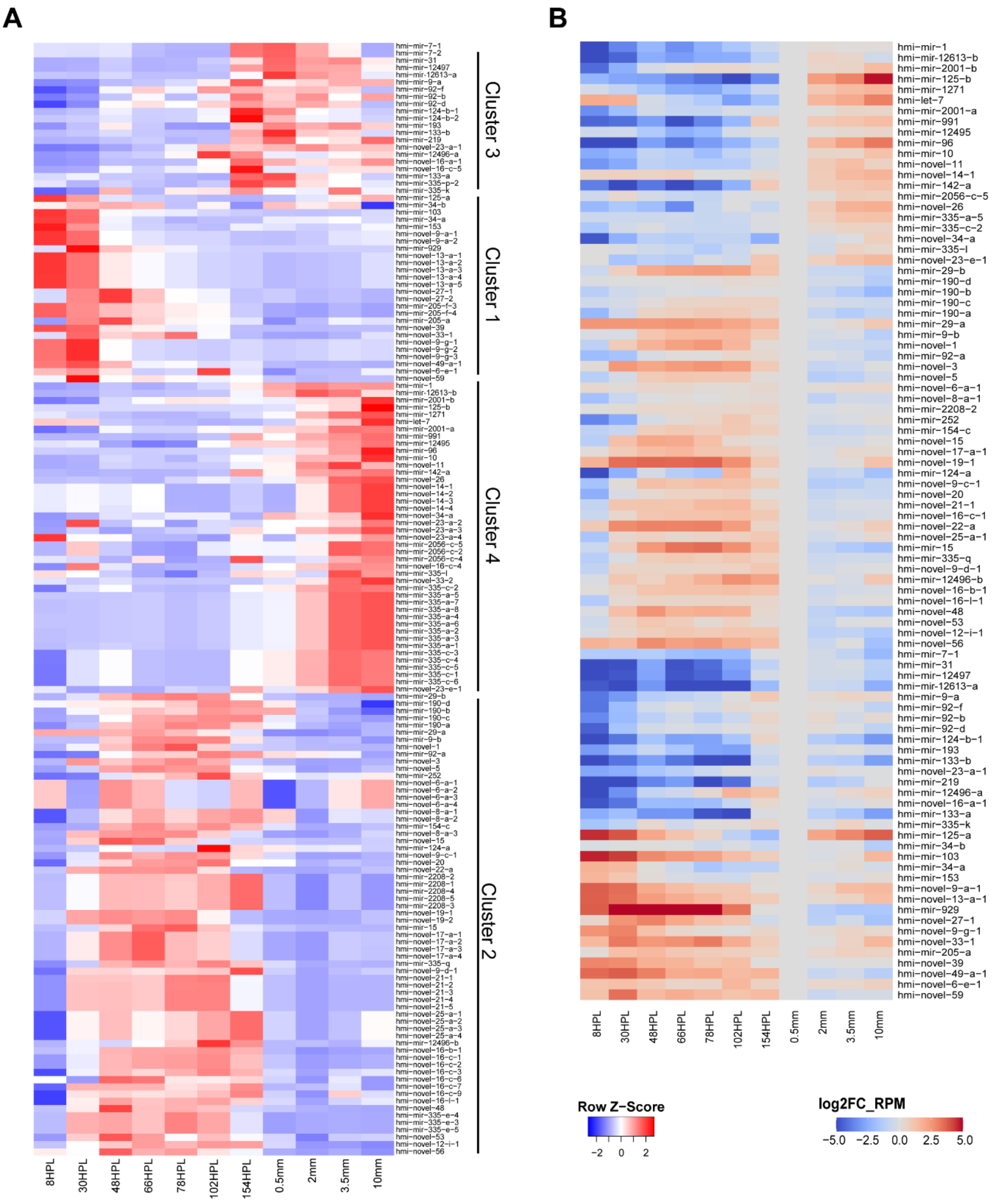


**Figure S8. Heatmaps of enrichment in expression (A) and fold change of normalized read counts (B) of *H. miamia* across the developmental stages.** The FC changes in RPM of each stage were normalized to that at the HL stage, which is the most synchronized stage with 16-hour windows.

Figure S9

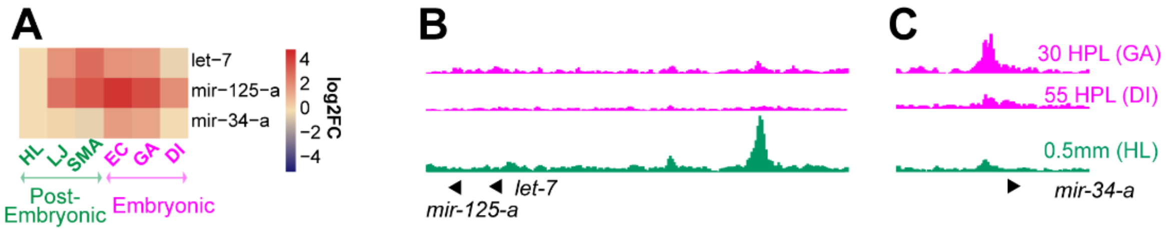


**Figure S9. Expression enrichment of *hmi-let-7* and *hmi-mir-125-a* during early embryonic development. A.** Heatmap showing the expression of *hmi-let-7, hmi-mir-125-a*, and *hmi-mir-34-a*. Expression is normalized to the HL (0.5 mm) stage. The post-embryonic developmental stages were positioned before the embryonic stages to emphasize the potential maternal deposition of miRNAs. **B-C.** Chromatin accessibility of the *let-7-mir-125-b* cluster locus (**B**) and the *mir-34-a* locus (**C**), as determined from previously published ATAC-seq data (5). The GA (30 HPL) and DI (55 HPL) time points represent early embryonic stages (green), and HL (0.5 mm) represents a post-embryonic stage (magenta) as data for later post-embryonic stages were unavailable. The displayed genomic regions span 10 kilobases.

Figure S10

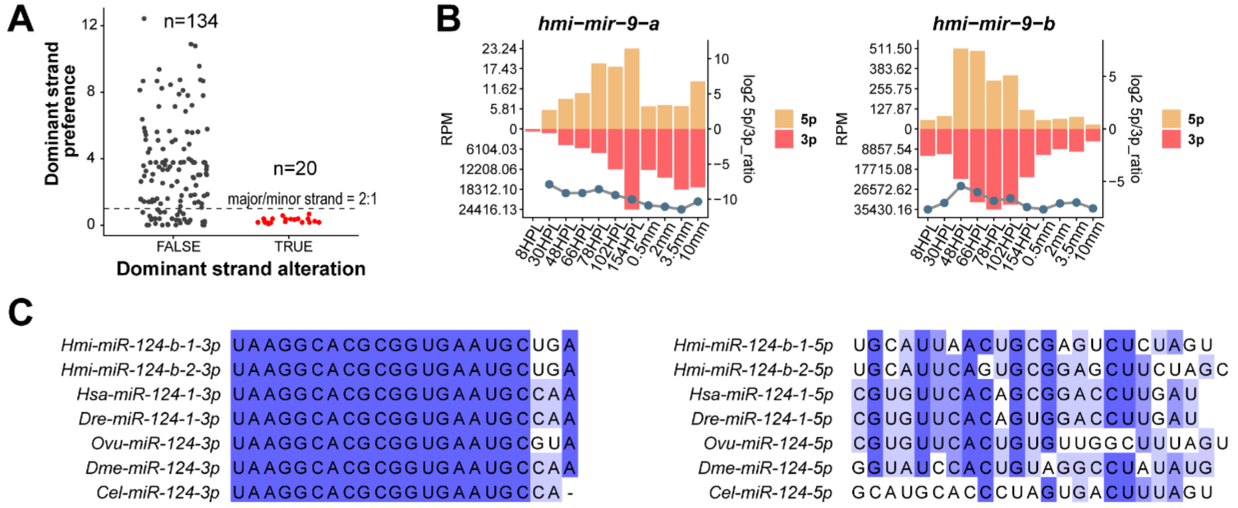


**Figure S10. Supplemental results for the *H. miamia* miRNA strand selection during development. A.** Minimum dominant strand preference across developmental stages for each highest-confidence miRNA. For each miRNA, dominant strand preference was quantified as the major-to-minor strand read-count ratio, and the minimum ratio observed across developmental stages is plotted. The dashed line indicates the arbitrary cutoff (major/minor = 2). Red points represent miRNAs that exhibit dominant strand alterations at any stage. **B.** Expression profiles of 5p and 3p strands and their strand preference. Bars indicate normalized read counts (RPM; left y-axis) of the 5p and 3p strands of *hmi-mir-9-a/b* , whereas points connected by lines indicate the 5p/3p ratio (right y-axis)**.** Note that unlike Figure 4, the y axis scales were rearranged to the RPM limits. **C.** Sequence alignment of the 3p and 5p strand of MIR-124 families.

Figure S11

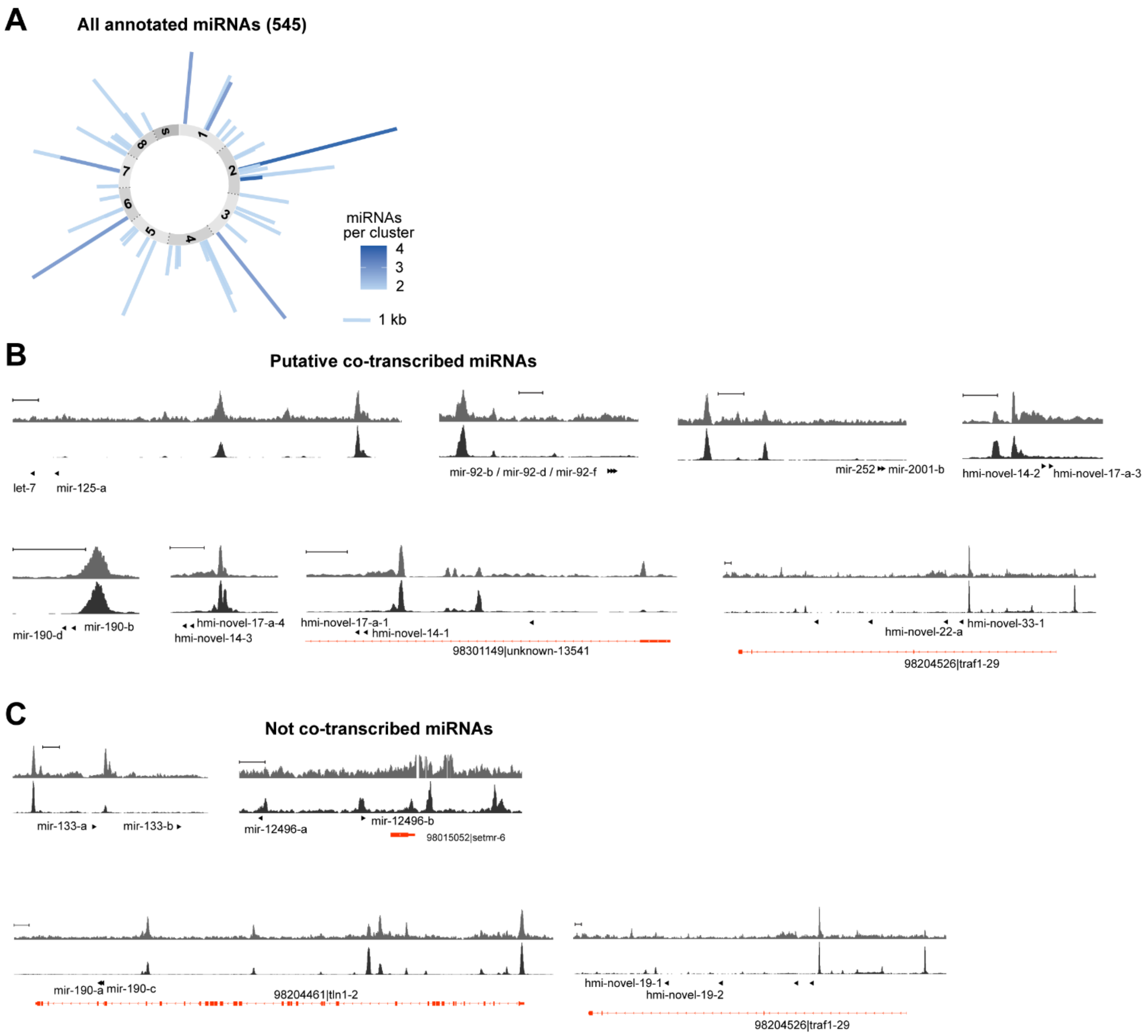


**Figure S11. Supplemental information on clustered miRNAs. A.** Summary of the genomic organization of clustered miRNAs among all 545 annotated miRNAs. Bar lengths indicate the genomic span of each cluster. **B-C.** Chromatin accessibility profiles of clustered miRNAs. For each panel, the upper tracks show the merged chromatin accessibility profiles from seven developmental stages from gastrula to hatchlings (5); the lower tracks show the merged chromatin accessibility profiles of the middle body region from late juveniles (6). Scale bars indicate 1 kb. Panel B shows clusters predicted to be co-transcribed based on shared putative promoters. In the *hmi-novel-17-a-1/hmi-novel-14-1/mir-335-c-3* cluster, although *hmi-novel-17-a-1* and *hmi-novel-14-1* are intronic, a putative promoter was predicted upstream and is not likely to be belong to the host gene, suggesting that these two miRNA genes may be co-transcribed in an independent primary transcript. In contrast, *mir-335-c-3* in that cluster was not considered as part of the co-transcribed unit. Panel C shows clusters predicted not to be co-transcribed because of independent putative promoters, intronic localization, or on opposite strands.

Figure S12

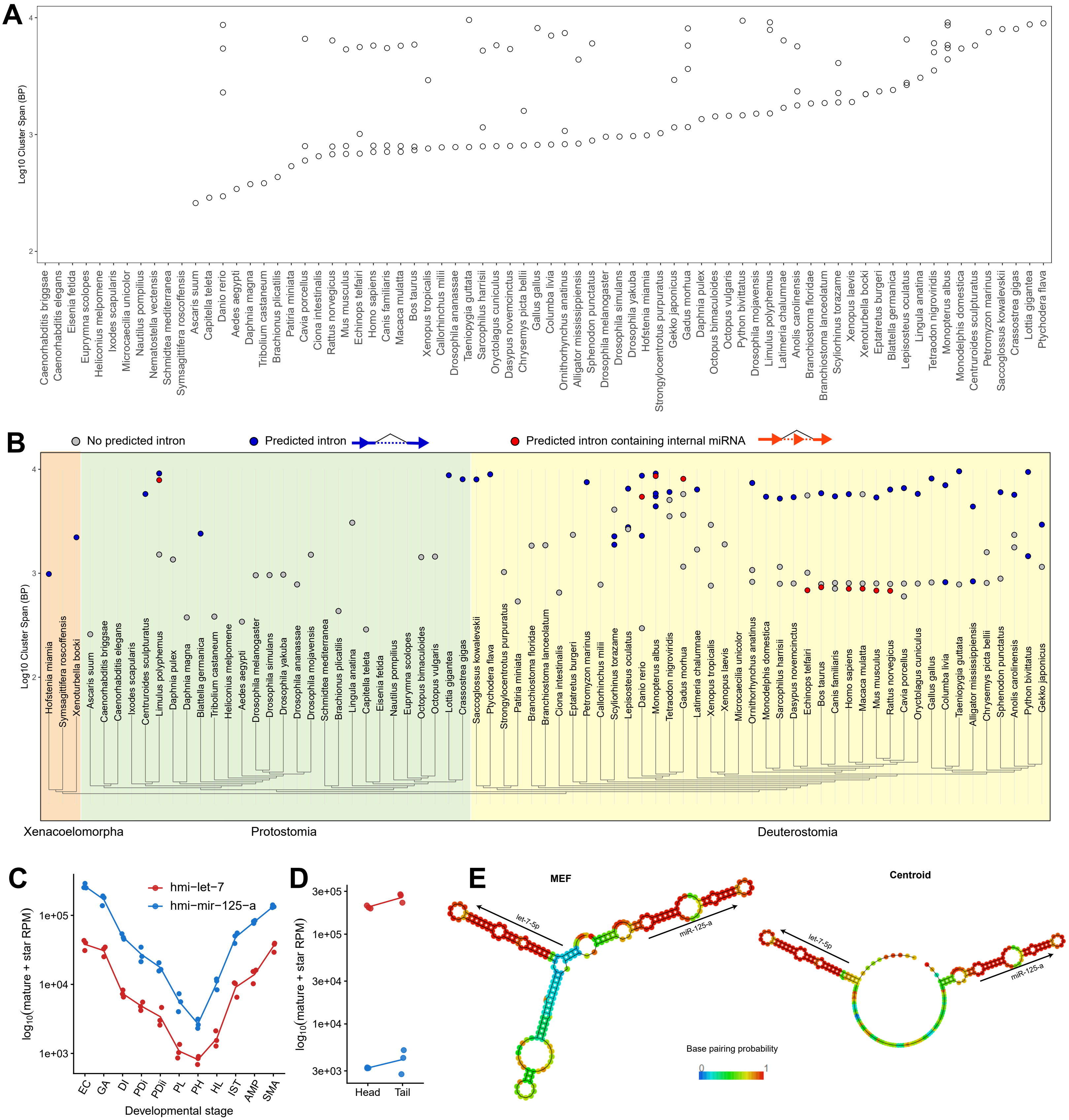

A
Log10 Cluster Span (BP)
2
3
4
Caenorhabditis briggsae
Caenorhabditis elegans
Eisenia fetida
Euprymna scolopes
Heliconius melpomene
Ixodes scapularis
Microcaecilia unicolor
Nautilus pompilius
Nematostella vectensis
Schmidtea mediterranea
Symsagittifera roscoffensis
Ascaris suum
Capitella teleta
Danio rerio
Aedes aegypti
Daphnia magna
Tribolium castaneum
Brachionus plicatilis
Patiria miniata
Cavia porcellus
Ciona intestinalis
Rattus norvegicus
Mus musculus
Echinops telfairi
Homo sapiens
Canis familiaris
Macaca mulatta
Bos taurus
Xenopus tropicalis
Callorhinchus milii
Drosophila ananassae
Taeniopygia guttata
Sarcophilus harrisii
Oryctolagus cuniculus
Dasypus novemcinctus
Chrysemys picta bellii
Gallus gallus
Columba livia
Ornithorhynchus anatinus
Alligator mississippiensis
Sphenodon punctatus
Drosophila melanogaster
Drosophila simulans
Drosophila yakuba
Hofstenia miamia
Strongylocentrotus purpuratus
Gekko japonicus
Gadus morhua
Daphnia pulex
Octopus bimaculoides
Octopus vulgaris
Python bivittatus
Drosophila mojavensis
Limulus polyphemus
Latimeria chalumnae
Anolis carolinensis
Branchiostoma floridae
Branchiostoma lanceolatum
Scyliorhinus torazame
Xenopus laevis
Xenoturbella bocki
Eptatretus burgeri
Blattella germanica
Lepisosteus oculatus
Lingula anatina
Tetraodon nigroviridis
Monopterus albus
Monodelphis domestica
Centruroides sculpturatus
Petromyzon marinus
Saccoglossus kowalevskii
Crassostrea gigas
Lottia gigantea
Ptychodera flava
B
No predicted intron
Predicted intron
Predicted intron containing internal miRNA
Log10 Cluster Span (BP)
2
3
4
Hofstenia miamia
Symsagittifera roscoffensis
Xenoturbella bocki
Ascaris suum
Caenorhabditis briggsae
Caenorhabditis elegans
Ixodes scapularis
Centruroides sculpturatus
Limulus polyphemus
Daphnia pulex
Daphnia magna
Blattella germanica
Tribolium castaneum
Heliconius melpomene
Aedes aegypti
Drosophila melanogaster
Drosophila simulans
Drosophila yakuba
Drosophila ananassae
Drosophila mojavensis
Schmidtea mediterranea
Brachionus plicatilis
Lingula anatina
Capitella teleta
Eisenia fetida
Nautilus pompilius
Euprymna scolopes
Octopus bimaculoides
Octopus vulgaris
Lottia gigantea
Crassostrea gigas
Saccoglossus kowalevskii
Ptychodera flava
Strongylocentrotus purpuratus
Patiria miniata
Branchiostoma floridae
Branchiostoma lanceolatum
Ciona intestinalis
Eptatretus burgeri
Petromyzon marinus
Callorhinchus milii
Scyliorhinus torazame
Lepisosteus oculatus
Danio rerio
Monopterus albus
Tetraodon nigroviridis
Gadus morhua
Latimeria chalumnae
Xenopus tropicalis
Xenopus laevis
Microcaecilia unicolor
Ornithorhynchus anatinus
Monodelphis domestica
Sarcophilus harrisii
Dasypus novemcinctus
Echinops telfairi
Bos taurus
Canis familiaris
Homo sapiens
Macaca mulatta
Mus musculus
Rattus norvegicus
Cavia porcellus
Oryctolagus cuniculus
Gallus gallus
Columba livia
Taeniopygia guttata
Alligator mississippiensis
Chrysemys picta bellii
Sphenodon punctatus
Anolis carolinensis
Python bivittatus
Gekko japonicus
Xenacoelomorpha
Protostomia
Deuterostomia
C
hmi-let-7
hmi-mir-125-a
log10(mature + star RPM)
1e+03
1e+04
1e+05
EC
GA
DI
PDi
PDii
PL
PH
HL
IST
AMP
SMA
Developmental stage
D
log10(mature + star RPM)
3e+03
1e+04
3e+04
1e+05
3e+05
Head
Tail
E
MEF
let-7-5p
miR-125-a
Centroid
let-7-5p
miR-125-a
Base pairing probability
0
1

**Figure S12. Supplemental data for the splicing of *hmi-let-7/hmi-mir-125-a*. A.** Distribution of genomic spans of clustered miRNA s of LET-7/MIR-125/MIR-100 families from the 73 bilaterian species containing members of LET-7/MIR-125/MIR-100 families. Each point represents a cluster spanning less than 10 kb. Species are ordered by the genomic span of their shortest cluster. **B.** Summary of cluster spans and predicted intron. The plot is formatted as in Figure 5E but uses the highest ASSP splice donor and acceptor score thresholds (7.0 for donor and 9.0 for acceptor; Figure 5E uses the default threshold of 2.2 and 4.5, respectively). **C-D.** Normalized expression (RPM) for *hmi-mir-125-a* and *hmi-let-7* during *H. miamia* development (C) and between head and tail fragments (D). **E.** Predicted secondary structure prediction of the spliced *let-7/mir-125-a* primary transcript generated using RNAfold with default settings. Both the minimum free energy (MFE; left) and centroid (right) structures are shown. Colors indicate base-pairing probability.

Figure S13

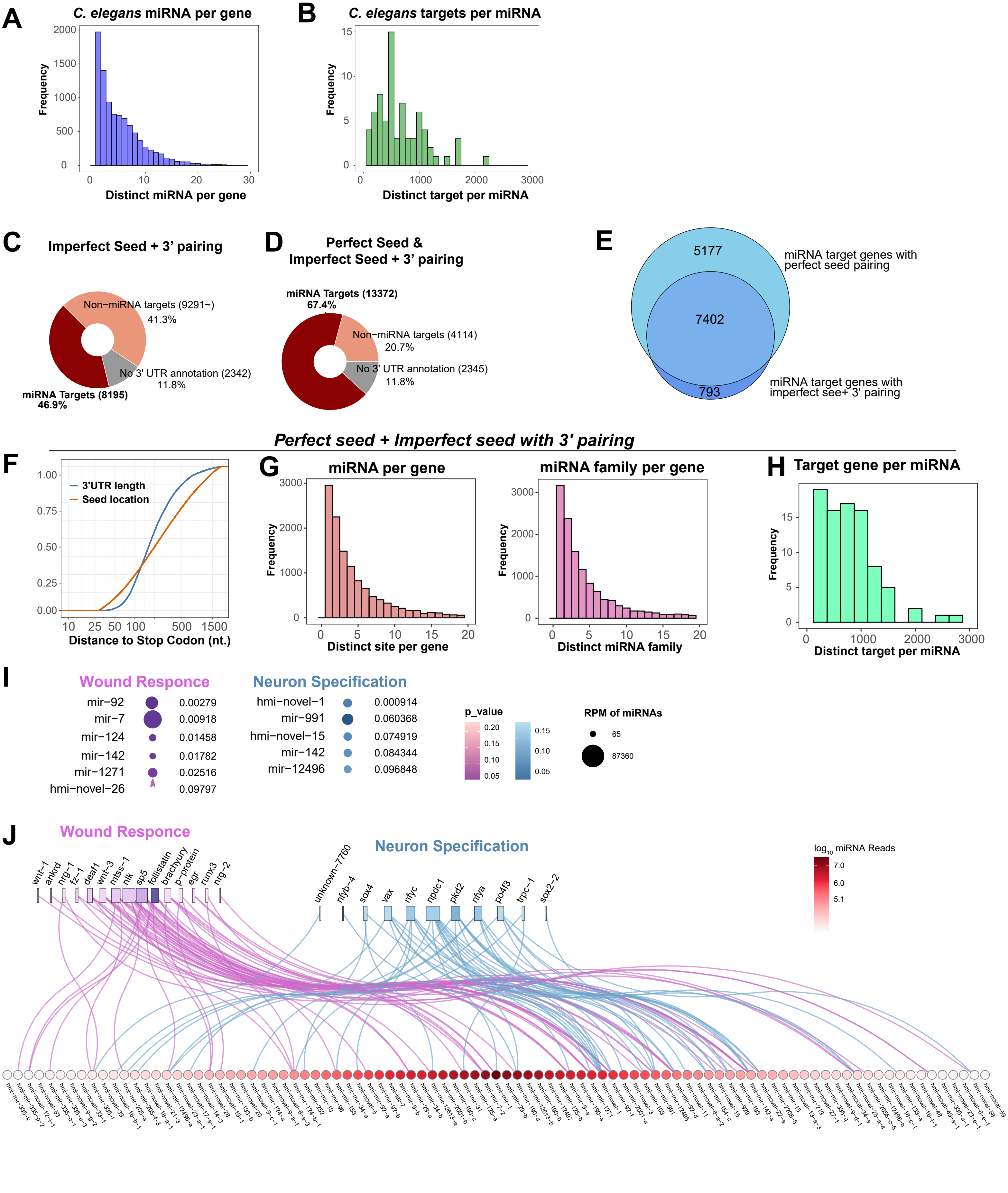
A
C. elegans miRNA per gene
Frequency
Distinct miRNA per gene
B
C. elegans targets per miRNA
Frequency
Distinct target per miRNA
C
Imperfect Seed + 3' pairing
Non−miRNA targets (9291~)
41.3%
No 3' UTR annotation (2342)
11.8%
miRNA Targets (8195)
46.9%
D
Perfect Seed &
Imperfect Seed + 3' pairing
miRNA Targets (13372)
67.4%
Non−miRNA targets (4114)
20.7%
No 3' UTR annotation (2345)
11.8%
E
5177
7402
793
miRNA target genes with perfect seed pairing
miRNA target genes with imperfect see+ 3' pairing
Perfect seed + Imperfect seed with 3' pairing
F
3'UTR length
Seed location
Distance to Stop Codon (nt.)
G
miRNA per gene
Frequency
Distinct site per gene
miRNA family per gene
Frequency
Distinct miRNA family
H
Target gene per miRNA
Frequency
Distinct target per miRNA
I
Wound Responce
mir−92 0.00279
mir−7 0.00918
mir−124 0.01458
mir−142 0.01782
mir−1271 0.02516
hmi−novel−26 0.09797
Neuron Specification
hmi−novel−1 0.000914
mir−991 0.060368
hmi−novel−15 0.074919
mir−142 0.084344
mir−12496 0.096848
p_value
RPM of miRNAs
65
87360
J
Wound Responce
wnt−1
ankrd
nrg−1
fz−1
deaf1
wnt−3
mtss−1
nlk
sp5
follistatin
brachyury
p−protein
egr
runx3
nrg−2
Neuron Specification
unknown−7760
nfyb−4
sox4
vax
nfyc
npdc1
pkd2
nfya
po4f3
trpc−1
sox2−2
log10 miRNA Reads
7.0
6.0
5.1

**Figure S13. Supplemental results for the *H. miamia* genome annotation and miRNA target analysis. A-B.** The average number of miRNAs that regulate a protein-coding gene (A) and the number of targets per miRNA (B) in *C. elegans*. **C.** Proportions of *H. miamia* protein-coding genes categorized by miRNA regulation potential. Only miRNA target sites with imperfect seed pairing plus 3' pairing are included in this analysis. For this analysis, imperfect seed pairing allows GU wobble pairs or a single-nucleotide bulge on the target strand at g5-g7 positions. The 3' pairing requires at least three consecutive Watson-Crick base pairs at g11-g16 (1). **D.** Proportions of *H. miamia* protein-coding genes categorized by their miRNA regulatory potential, including both perfect seed pairing configurations (Figure 6D) and imperfect seed pairing plus 3' pairing configurations (Figure S10B). **E.** Venn diagram showing the overlap of protein-coding genes containing miRNA targets sites with perfect seed pairing configuration and with imperfect seed + 3' pairing configuration. **F.** Empirical cumulative distribution of the distance from stop codon comparing the 3' UTR lengths and miRNA seed location (g2). **G.** Distribution of the number of predicted miRNA target sites per gene (left panel) and the number of miRNA seed families per gene (right panel). **H.** Distribution of the numbers of predicted targets per miRNA. In **F-H**, both perfect seed pairing configuration and imperfect seed + 3' pairing configuration were included for comparison with Figure 6E-G. **I.** Enrichment for target sites of each miRNA seed family in the wound response network (magenta) and the neural differentiation network (blue). The p-values indicate the cumulative probability of a one-tailed hypergeometric test. The miRNA families that significantly enriched for the two gene groups with $p < 0.1$ are shown. **I-J.** Enrichment for targets sites of each miRNA (**I**) and putative miRNA-target interactome (**J**) using both perfect seed pairing and the imperfect seed pairing plus 3' pairing configuration. Threshold and analysis process are identical to Figure 6J-K. Because this interactome incorporates 3' pairing, individual miRNAs are shown instead of being grouped by seed families.
